# FaRIF at the Core of Strawberry Fruit Ripening: Deciphering its Targets and Interaction Networks

**DOI:** 10.1101/2025.01.13.631932

**Authors:** Carmen Martín-Pizarro, José M. Franco-Zorrilla, María Florencia Perotti, Rosa Lozano-Durán, Guozheng Qin, David Posé

**Author notes:** Corresponding authors: Carmen Martín-Pizarro, David Posé. The authors responsible for distribution of materials integral to the findings presented in this article are: Carmen Martín Pizarro and David Posé.

## Abstract

Ripening Inducing Factor (RIF) is a key NAC transcription factor regulating strawberry fruit ripening. Previous studies using *RIF-*RNAi and overexpression lines in *Fragaria × ananassa* and CRISPR knock-out lines in *F. vesca* have established the role of RIF in controlling ABA biosynthesis and signaling, cell wall remodeling, and secondary metabolism.

In this study, we deciphered FaRIF’s transcriptional regulatory network by combining ChIP-seq-based identification of its direct targets with an analysis of *FaRIF*-RNAi transcriptome data. These analyses revealed FaRIF’s direct role in multiple aspects of strawberry fruit ripening, including the regulation of ripening-related transcription factors, phytohormone content and signaling, primary and secondary metabolism, and cell wall degradation. Additionally, using the TurboID-based proximity labeling approach, we have identified FaRIF interactors, including proteins involved in mRNA and protein homeostasis, as well as several NAC transcription factors. Among these, FaNAC034 was found to synergistically enhance FaRIF’s transcriptional activity.

This integrative analysis, combining transcriptome analysis, *in vivo* ChIP-seq, and proximity labeling, broadens our knowledge of FaRIF-mediated transcriptional networks and interaction partners, providing valuable insights into the molecular mechanisms underlying strawberry fruit ripening regulation by this transcription factor.

## Introduction

Cultivated strawberry (*Fragaria × ananassa* Duch.) is one of the most economically significant berry crops worldwide (Simpson, 2018), primarily due to its unique organoleptic attributes and nutritional value (Azodanlou et al., 2003; Ikegaya et al., 2019; Newerli-Guz et al., 2023). These characteristics develop during fruit ripening, a highly coordinated developmental process that has evolved to facilitate seed dispersal, an essential factor in evolutionary success (Klee and Giovannoni, 2011). Ripening is characterized by the softening of the fruit and an increase in the sugars, flavor-enhancing volatiles, and essential micronutrients content. The mechanisms underlying these processes primarily rely on the action of phytohormones (Perotti et al., 2023), epigenetic modifications (Li et al., 2022b; Pan et al., 2024), and transcription factors (TFs) (Sánchez-Gómez et al., 2022).

As a non-climacteric fruit, strawberry ripening is mainly regulated by abscisic acid (ABA) (Chai et al., 2011; Jia et al., 2011; Li et al., 2022a), whose levels increase during fruit ripening (Chen et al., 2011; Symons et al., 2012; Gu et al., 2019). In addition to ABA, other phytohormones also contribute to strawberry fruit development and ripening (Perotti et al., 2023). Notably, auxins and gibberellins play pivotal roles during early developmental stages, inhibiting ripening by suppressing ABA accumulation through the activation of *CYP707A4a* expression, a cytochrome P450 monooxygenase involved in ABA catabolism (Liao et al., 2018). Epigenetic modifications, including changes in DNA methylation (Cheng et al., 2018; Zhou et al., 2021; Martínez-Rivas et al., 2022) and histone modifications (Pan et al., 2024), have also been reported to influence strawberry ripening, further underscoring their significance in this process.

In recent years, a number of TFs have been identified as important regulators of ripening, most of which have a specific role in a particular ripening-related process (Sánchez-Gómez et al., 2022). Among these, MYB10, a member of the R2R3-type MYB TF family, is a key regulator of anthocyanin biosynthesis and probably the most extensively studied TF in strawberry (Lin-Wang et al., 2010; Medina-Puche et al., 2013; Lin-Wang et al., 2014; Hawkins et al., 2016; Zhang et al., 2017; Castillejo et al., 2020; Manivannan et al., 2021). However, despite significant progress in understanding the role of TFs in strawberry fruit ripening regulation, the transcriptional regulatory networks they control and the mechanisms regulating their activity still remain poorly characterized.

NAC (NAM, ATAF, and CUC) TFs constitute a large plant-specific family involved in diverse developmental processes and responses to environmental stimuli (Olsen et al., 2005), with a number of them being involved in fruit ripening in different species (Forlani et al., 2021; Liu et al., 2022a). These proteins are characterized by a conserved N-terminal region, known as the NAC domain, responsible for DNA recognition, binding, and dimerization, and a variable C-terminal region that defines the different NACs subgroups. In the woodland strawberry *F. vesca*, one of the diploid ancestor species of *F. × ananassa* (Edger et al., 2019), 112 *NAC* genes have been identified (Moyano et al., 2018), although only two have been studied in detail. One of these, *FcNAC1*, the ortholog of *NAC022* in the beach strawberry (*F. chiloensis*), was shown to activate the expression of *FcPL*, a gene encoding the cell-wall remodeling enzyme pectate lyase (Carrasco-Orellana et al., 2018). However, the only functional studies for a strawberry NAC TF have been focused on *NAC035*, named *Ripening Inducing Factor* (*RIF*) in both *F. × ananassa* (*FaRIF*) and *F. vesca* (*FvRIF*) (Martín-Pizarro et al., 2021; Li et al., 2023). In these studies, stable RNAi-silencing and CRISPR knockout lines were established in each species, along with overexpression lines in both. Transcriptomic and metabolomic profiling, combined with phenotypic characterization in *F. × ananassa* lines with altered *FaRIF* expression, revealed that this TF regulates several ripening-associated processes, including cell wall degradation, anthocyanin biosynthesis, and the accumulation of sugars, organic acids, and volatiles. It also influences the aerobic/anaerobic metabolic balance, a key determinant in the onset of strawberry fruit ripening (Wang et al., 2017). Notably, FaRIF was shown to regulate ABA accumulation, suggesting an upstream regulatory role for this TF (Martín-Pizarro et al., 2021). The characterization of its ortholog, *FvRIF*, in *F. vesca* further confirmed its central regulatory role of this TF, displaying the *Fvrif* knockout mutant lines a complete blockage of the ripening process (Li et al., 2023). In this study, FvRIF was shown to interact with and serves as a substrate for MAP kinase 6 (FvMAPK6), being the phosphorylation at Thr-310 essential for its transcriptional activity. Additionally, the DNA binding sites of FvRIF using DNA affinity purification sequencing (DAP-seq) were identified, revealing several structural genes involved in the anthocyanin pathway, cell-wall degradation, sugar metabolism, and aroma compounds biosynthesis among its direct target genes.

DAP-seq is a powerful, fast, and cost-effective technique for identifying DNA binding sites of TFs or other DNA-associated proteins (Bartlett et al., 2017). To date, and in addition to the study of FvRIF, only a few recent studies have employed this approach to identify TF target genes in strawberry, including the APETALA2 (AP2) TF BARE RECEPTACLE (BRE) (Lu et al., 2024b) and FvTCP7 (Chen et al., 2024), which are involved in strawberry floral organogenesis and fruit ripening, respectively. However, although of great value, DAP-seq, as an *in vitro* approach, lacks the chromatin context of *in vivo* interactions and does not account for the influence of other protein interactors or cofactors on TF binding sites, which can be captured using Chromatin Immunoprecipitation (ChIP) followed by deep sequencing (ChIP-seq). Probably due to the extremely recalcitrant nature of strawberry, ChIP has only been applied, and only followed by qPCR validation, in a few studies in strawberry to validate target genes for FvWRKY48 (Zhang et al., 2022b), BRE (Lu et al., 2024b), and FvRIF itself (Li et al., 2023). Notably, only a few examples of ChIP-seq application have been reported in this species, focusing on the study of chromatin states through histone modification analysis (Huang et al., 2020; Luo et al., 2022; Pan et al., 2024; Baldwin et al., 2024), while no studies to date have addressed the identification of TF target genes.

In this work, we have advanced our understanding of FaRIF role in the regulation of strawberry fruit ripening. First, we have set up a protocol for successfully perform a ChIP-seq experiment on a TF and we have identified its target genes *in vivo* in *F. × ananassa.* By integrating these results with the analysis of a transcriptome dataset from *RIF*-RNAi lines, mapped to the latest octoploid reference genome annotation of *F. × ananassa* cv Camarosa (Liu et al., 2021), we uncovered the gene regulatory network governed by this TF, both directly and indirectly.

Additionally, we optimized and applied the TurboID-based proximity labeling approach in strawberry fruits to investigate FaRIF’s proximal interactome (i.e. proxitome) *in vivo.* This analysis revealed a diverse set of putative FaRIF interactors, including other NAC family members, such as the ripening-related FaNAC034 and FaNAC021, as well as proteins involved in processes like protein folding and mRNA stability. In summary, this study provides valuable insights into the molecular mechanisms underpinning the complex ripening process in strawberry mediated by this key TF. Furthermore, it offers detailed methodologies that will benefit researchers interested in applying these omics approaches to a complex species like strawberry.

## Results

### Analysis of FaRIF protein homoeologs and characterization of their expression profile in *F.* × ananassa cv. Camarosa

*FaRIF*, whose ortholog in *F. vesca* (*FvRIF*) is encoded by the gene FvH4_3g20700, is encoded by four homoeologs in the *F.* × ananassa cv. Camarosa genome (Edger et al., 2019; Liu et al., 2021), located on chromosomes 3A (*FaRIF(3A)*, FxaC_9g32650 or maker-Fvb3-4-augustus-gene-182.31), 3B (*FaRIF(3B)*, FxaC_10g22240 or augustus_masked-Fvb3-2-processed-gene-121.1), 3C (*FaRIF(3C)*, FxaC_11g20020 or maker-Fvb3-3-augustus-gene-106.29), and 3D (*FaRIF(3D)*, FxaC_12g28600 or maker-Fvb3-1-augustus-gene-197.26) (Supplementary Data Set 1), according to the chromosome nomenclature suggested by Hardigan and collaborators (Hardigan et al., 2021). Protein sequence alignment of the four homoeologs revealed that FaRIF(3A), (3B) and (3C) shared most residues with FvRIF, with FaRIF(3A) exhibiting an identical protein sequence to the latter (Supplementary Fig. S1). However, *FaRIF(3D)* encoded for a shorter protein, lacking the first 144 residues of the conserved N-terminal region and consequently missing most of the NAC domain (Supplementary Fig. S1). A protein structure prediction using AlphaFold 3 (Abramson et al., 2024) (Fig. 1A, Supplementary Fig. S2), indicated the absence in FaRIF(3D) of essential residues needed for DNA binding and NAC protein dimerization (Ernst et al., 2004), suggesting that this homoeolog is nonfunctional. Based on the reanalysis of the transcriptome changes during *F.* × ananassa cv. Camarosa ripening (Liu et al., 2021), and as previously reported for *FvRIF* (Li et al., 2023) and for *FaRIF* using the *F. vesca* as the reference genome (Martín-Pizarro et al., 2021), *FaRIF* expression increased during the ripening process of receptacles (Fig. 1B). This expression pattern was observed for all *FaRIF* homoeologs, except for the truncated homoeolog *FaRIF(3D)*, whose expression remained low throughout the process (Fig. 1B). Thus, these findings support that the homoeolog in this subgenome does not contribute to the regulation of strawberry ripening. Furthermore, *FaRIF(3A)*, the identical ortholog to *FvRIF* located in the *F. vesc*a’s subgenome, was found to be the most highly expressed *FaRIF* homoeolog (Fig. 1B).

**Figure 1.**
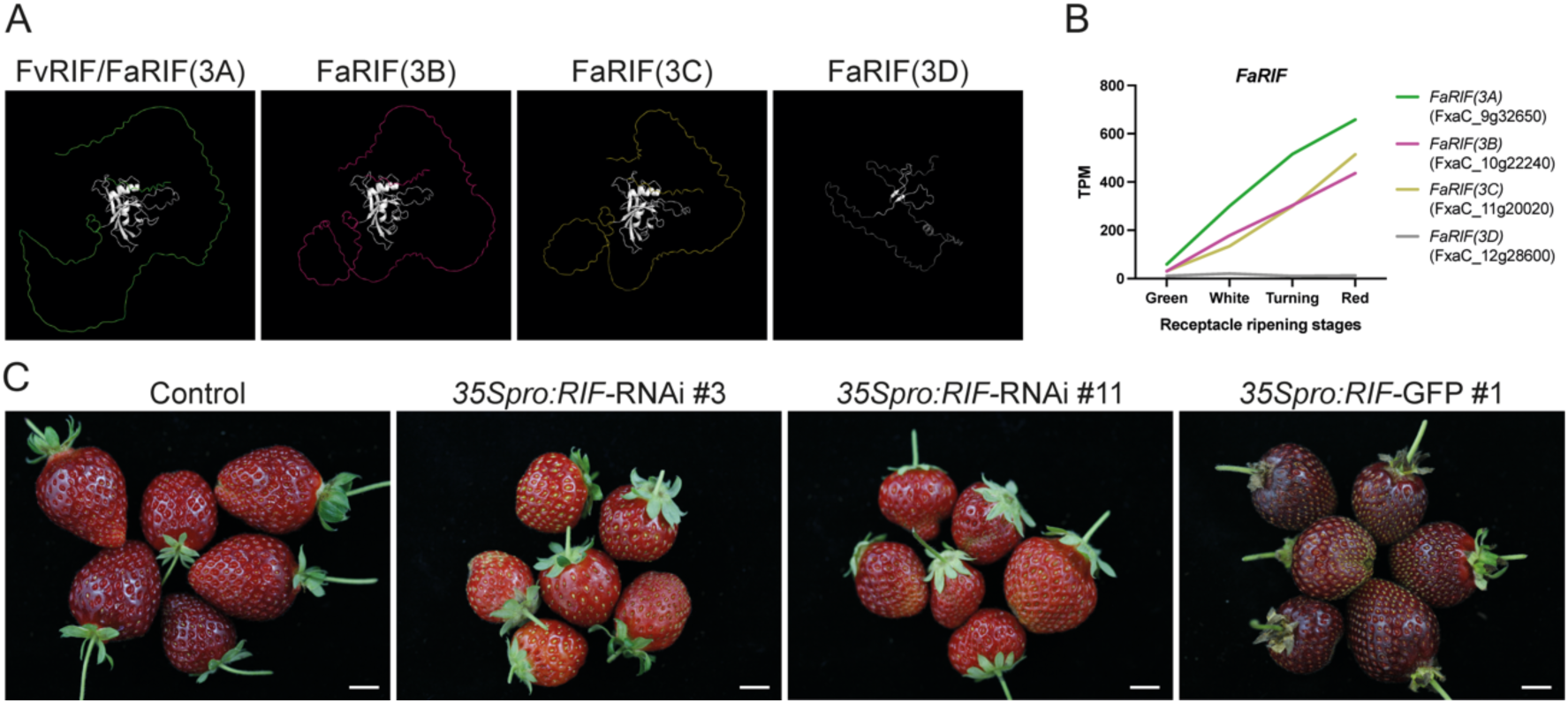
Protein structures, gene expression during strawberry fruit ripening and phenotypes of *FaRIF* silencing and overexpression lines. **A)** Predicted protein structures of FvRIF and FaRIFs homoeolog proteins using Alphafold 3 (Abramson et al., 2024). The predicted models were aligned to model 0 of the FvRIF/FaRIF(3A) protein using PyMol. The NAC domain is shown in white. **B)** Expression patterns of the four *FaRIF* homoeologs in *F.* × *ananassa* at four ripening stages in receptacles. Data from (Sánchez-Sevilla et al., 2017; Liu et al., 2021). Colors denote the four subgenomes encoding each *FaRIF* homoeolog: green (*F. vesca*), purple (*F. iinumae*), yellow (*F. nipponica*), and grey (*F. viridis*). **C)** Fruit phenotype at the red stage in control and stable *35Spro:RIF-*RNAi and *35Spro:RIF-GFP* transgenic lines. Scale bars = 1 cm.

### Identification of genome-wide *in vivo* FaRIF binding sites

Using our previously established GFP-tagged *FaRIF* overexpression line in *F.* × ananassa cv. Camarosa (Fig. 1C) (Martín-Pizarro et al., 2021), we aimed to elucidate the *in vivo* binding sites of FaRIF through a ChIP-seq assay. We processed two independent biological replicates of the *35Spro:RIF-GFP* #1 line, collecting both immunoprecipitated (IP) and DNA input samples. To validate the ChIP assay, we analyzed the enrichment of two known direct targets of FvRIF (*F. vesca*’s ortholog) (Li et al., 2023): *NAC042,* a ripening-induced NAC TF (Martín-Pizarro et al., 2021), and *PECTATE LYASE 2* (*PL2*), involved in cell-wall disassembly (Jiménez-Bermúdez et al., 2002), along with *FaRIF* itself. As a negative control, we selected a locus not expected to bind FaRIF (the first exon of FxaC_21g51230, a gene not expressed in fruits (Liu et al., 2021)). As shown in Fig. 2A, the two biological replicates of the FaRIF-GFP samples exhibited at least 4-fold enrichment for the three expected target genes, in contrast to the negative control. This validated the ChIP assay and supported the binding of both FvRIF and FaRIF to the promoters of *NAC042* and *PL2*, as well as to its own promoter, suggesting that FaRIF may control its own expression through feedback regulation.

**Figure 2.**
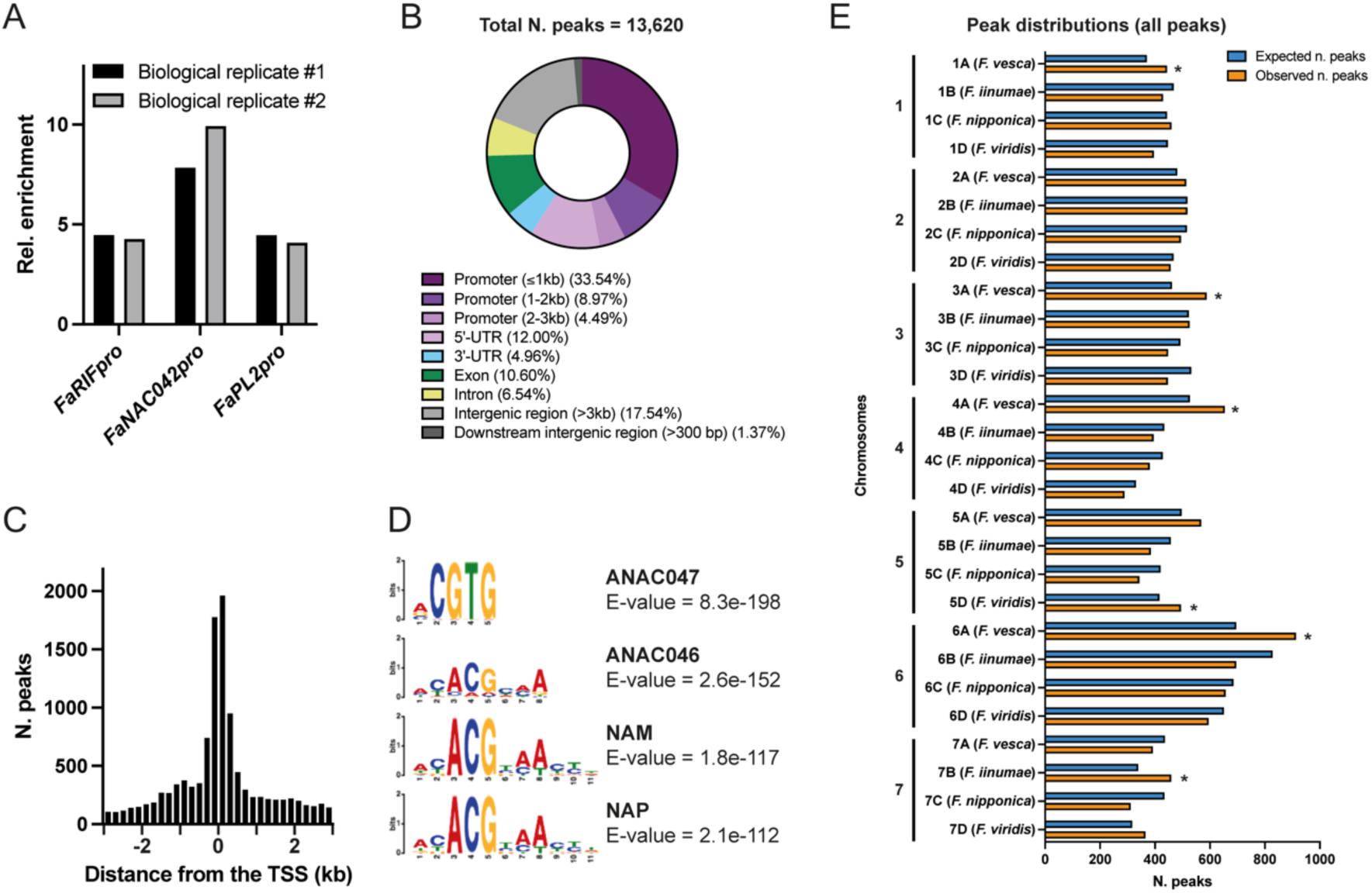
Genome-wide identification of FaRIF binding sites through ChIP-seq. **A)** ChIP-qPCR assay showing the enrichment of binding of FaRIF-GFP to the promoters of *FaRIF*, *FaNAC042*, and *FaPL2*, relative to the negative control (FxaC_21g51230). **B)** Distribution of FaRIF binding sites across genome features. **C)** Distribution of FaRIF binding sites 3 kb up- and downstream of the TSS. **D)** Most enriched NAC DNA logos for FaRIF DNA binding sites, corresponding to the mentioned Arabidopsis genes. **E)** Number of observed (orange bars) and expected (blue bars) binding sites for all peaks consistently identified in both biological replicates. The expected number of binding sites was calculated by dividing the total number of peaks for each chromosome set by the total size of that set. Asterisks (*) indicate statistically significant differences between observed and expected binding sites.

Subsequently, libraries were prepared for the two biological replicates of RIF-GFP IP and input samples and analyzed by high-throughput sequencing. A total of 13,620 peaks were identified across both biological replicates, therefore considered high-confidence binding sites for FaRIF (Supplementary Data Set 2). A peak distribution analysis showed that most binding sites were located within promoter regions, with 59% positioned within 3 kb upstream of the transcription start site (TSS), primarily 1 kb upstream (Fig. 2B). Additionally, examining a 3 kb region surrounding the TSS revealed that most peaks were centered on the TSS (Fig. 2C). A MEME analysis further identified significant enrichment of known NAC recognition motifs among the FaRIF binding sites, including those corresponding to ANAC047, ANAC046, NAM (the closest Arabidopsis homolog to FaRIF), and NAP (Fig. 2D), supporting the effectiveness of the ChIP-seq experiment.

Next, we examined whether FaRIF binding sites show positional bias in the genome by analyzing peak distribution across the homoeologous chromosomes of the four parental subgenomes of *F.* × ananassa, within the seven sets of chromosomes. While no preference was observed for chromosome 2, all other chromosomes exhibited higher representation of FaRIF binding sites in specific subgenomes. Notably, the *F. vesca* subgenomes contained more FaRIF binding sites on chromosomes 1, 3, 4, and 6 compared to the other subgenomes (Fig. 3E and Supplementary Data Set S3). Focusing on peaks located 2 kb upstream to 100 bp downstream of the TSS, regions most likely influencing gene expression regulation, we observed a similar FaRIF preference, except for chromosome 1, which showed no bias (Supplementary Fig. S3 and Data Set S3). Thus, these findings also align with the subgenome dominance of *F. vesca* reported by Edger and collaborators (Edger et al., 2019).

**Figure 3.**
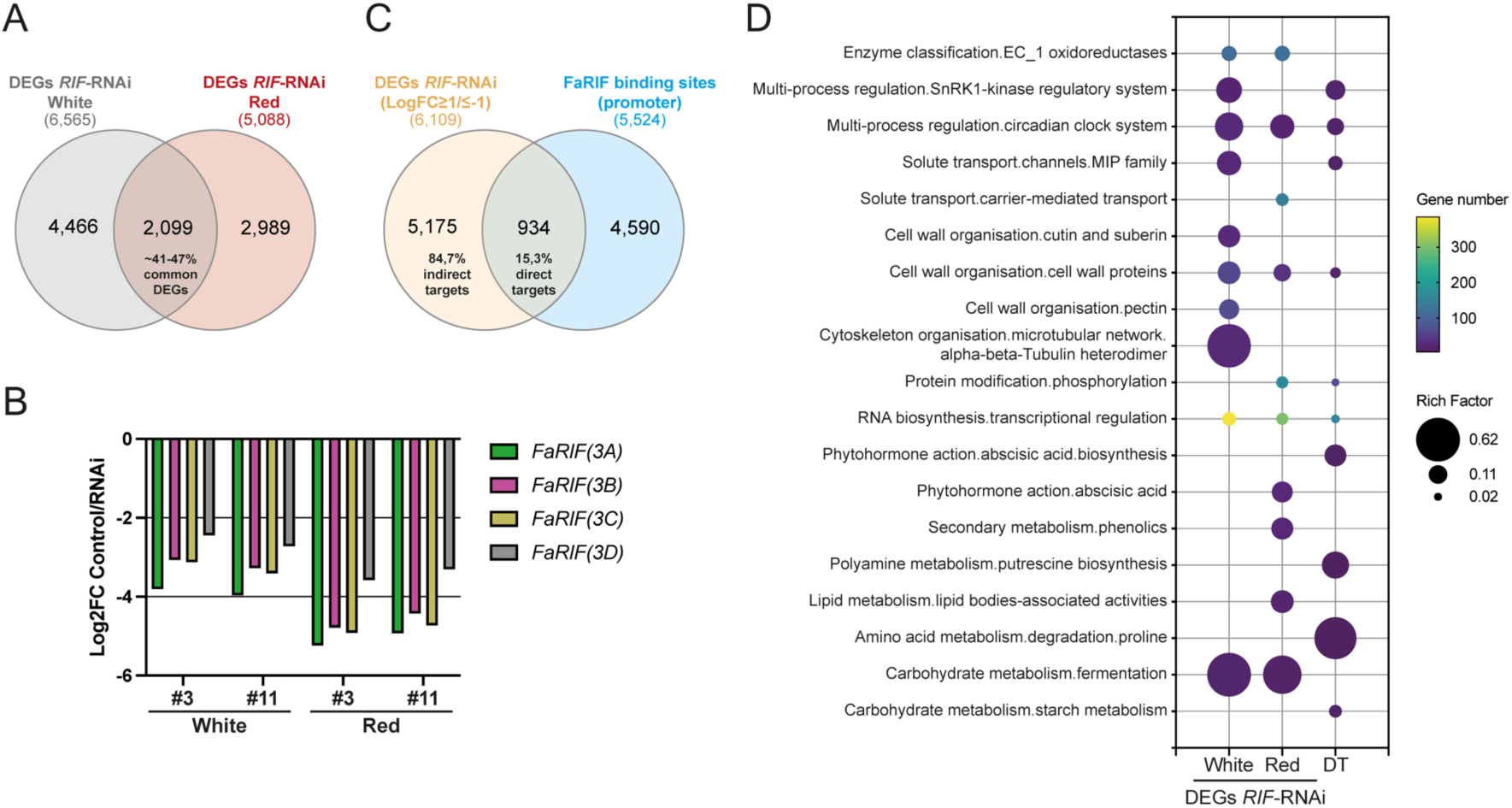
RNA-seq and ChIP-seq data reveal biological processes directly and indirectly regulated by FaRIF. **A)** Comparison of DEGs in receptacles of both independent *FaRIF-*RNAi lines (*35Spro:RIF-*RNAi #3 and #11) at the white and red ripening stages. **B)** Expression of the four *FaRIF* homoeologs (Log2FC control/RNAi ratio) in receptacles of both independent *FaRIF-*RNAi lines at the white and red ripening stages. **C)** Comparison of all DEGs identified in *FaRIF*-RNAi lines at both ripening stages and genes with FaRIF binding sites in the promoter regions. **D)** MapMan enrichment analysis of DEGs in *FaRIF-*RNAi receptacles (≥2-fold up- or downregulation and FDR ≤ 0,05 in the RNAi lines) at white and red stages and of FaRIF direct targets (DT). Top ten most statistically significant MapMan second-level bins levels for each analysis were selected for representation. Unique lower-level bins found within the top ten were also included.

### Identification of genes directly and indirectly regulated by FaRIF

To identify genes transcriptionally regulated by FaRIF, we analyzed previously generated RNA-seq data from receptacles at white and red stages of control and the two RNAi lines (*35Spro:RIF-*RNAi #3 and #11) (Fig. 1C). These datasets, originally mapped to the *F. vesca* reference genome (Martín-Pizarro et al., 2021), were re-mapped in this study to the *F.* × *ananassa* cv. Camarosa genome assembly v1.0.a1 (Edger et al., 2019) using the v1.0.a2 annotation version (Liu et al., 2021). Of the 108,447 annotated genes in the *F.* × *ananassa* genome, 6,565 and 5,088 genes were found differentially expressed (DEGs; false discovery rate (FDR) *P-value* correction ≤0.05) in both *FaRIF*-RNAi lines at the white and red stages, respectively, with 2,099 of them shared between the two RNAi lines (Fig. 3A, Supplementary Data Set 4). Furthermore, approximately 50% of the DEGs were either up- or downregulated at each ripening stage (Supplementary Data Set 4). Confirming the efficiency of the RNAi-mediated silencing, transcript levels of the four *FaRIF* homoeologs were significantly reduced in both transgenic lines and at both ripening stages (Fig. 3B, Supplementary Data Set 4).

Then, to identify genes directly or indirectly regulated by FaRIF, we compared the RNA-seq and ChIP-seq datasets under stringent criteria. Specifically, we selected those DEGs with at least a two-fold change (down- or upregulated; Log_2_(FC) ≤ -1 or ≥ 1; FDR ≤ 0.05) at the white and red stages (6,109 in total) in *FaRIF-*RNAi receptacles, and compared them to genes containing FaRIF binding sites within their promoter regions (from 2 kb upstream to 100 bp downstream of the TSSs; 5,524 genes). Overall, 15.3% of DEGs (934 *F.* × *ananassa* genes, corresponding to 718 unique *F. vesca* homoeologs) contained FaRIF binding sites, categorizing them as direct FaRIF targets (Fig. 3C; Supplementary Data Set S5). The remaining 84.7% of DEGs (5,175 *F.* × *ananassa* genes, corresponding to 3,484 unique *F. vesca* homoeologs) were therefore considered indirect targets. A stage-specific analysis identified 426 FaRIF-direct targets among the DEGs at the white stage, 308 at the red stage, and 200 shared between both stages (Supplementary Fig. S4A). Furthermore, of these direct targets, 691 genes (60.1%) were downregulated, while 443 (39%) were upregulated across both ripening stages (Supplementary Data Set S5), suggesting that FaRIF primarily functions as a transcriptional activator.

We next conducted a MapMan analysis to identify enriched functional categories among DEGs with more than two-fold change in expression in the *FaRIF*-RNAi lines at both ripening stages, as well as among the identified FaRIF direct targets (Schwacke et al., 2019) (Fig. 3D, Supplementary Data Set 6). At both stages, enriched processes included transcriptional regulation, enzyme activity, cell wall organization, and fermentation. At the red stage specifically, we observed significant enrichment in categories related to ABA and phenolic compounds, among others, supporting the role of FaRIF in directly or indirectly controlling key ripening-related processes. Interestingly, the analysis of FaRIF-direct targets further revealed that FaRIF directly regulates genes involved in transcriptional regulation, cell wall organization, and ABA biosynthesis, among others, highlighting its essential role in directly modulating these ripening-related processes.

### FaRIF regulates transcriptional cascades involved in strawberry ripening regulation

Within the Mapman bin related to transcriptional regulation, several TFs with roles in fruit development and/or ripening were found (Supplementary Data Set S7). First, *FaRIF*—specifically the most expressed homoeolog, *FaRIF(3A)*—was found among the direct targets (Fig. 4A), as previously confirmed by ChIP-qPCR, suggesting a positive feedback regulation of its own expression. Additionally, homoeologs of five other *NAC* genes induced during receptacle ripening, including *FaNAC04*, *FaNAC06* (both FaRIF-direct targets), *FaNAC022*, *FaNAC033*, and *FaNAC042*, were differentially expressed in at least one ripening stage (Fig. 4C, Supplementary Fig. S5A). Notably, all three *FaNAC042* homoeologs were consistently downregulated at both ripening stages in the two RNAi lines (Supplementary Fig. S5A), with one also identified as a FaRIF-direct target (Fig. 4A, C). A transactivation assay in *N. benthamiana* leaves confirmed FaRIF-mediated activation of its own promoter and that of *FaNAC042*, supporting FaRIF’s direct regulatory role over both genes (Fig. 4B).

**Figure 4.**
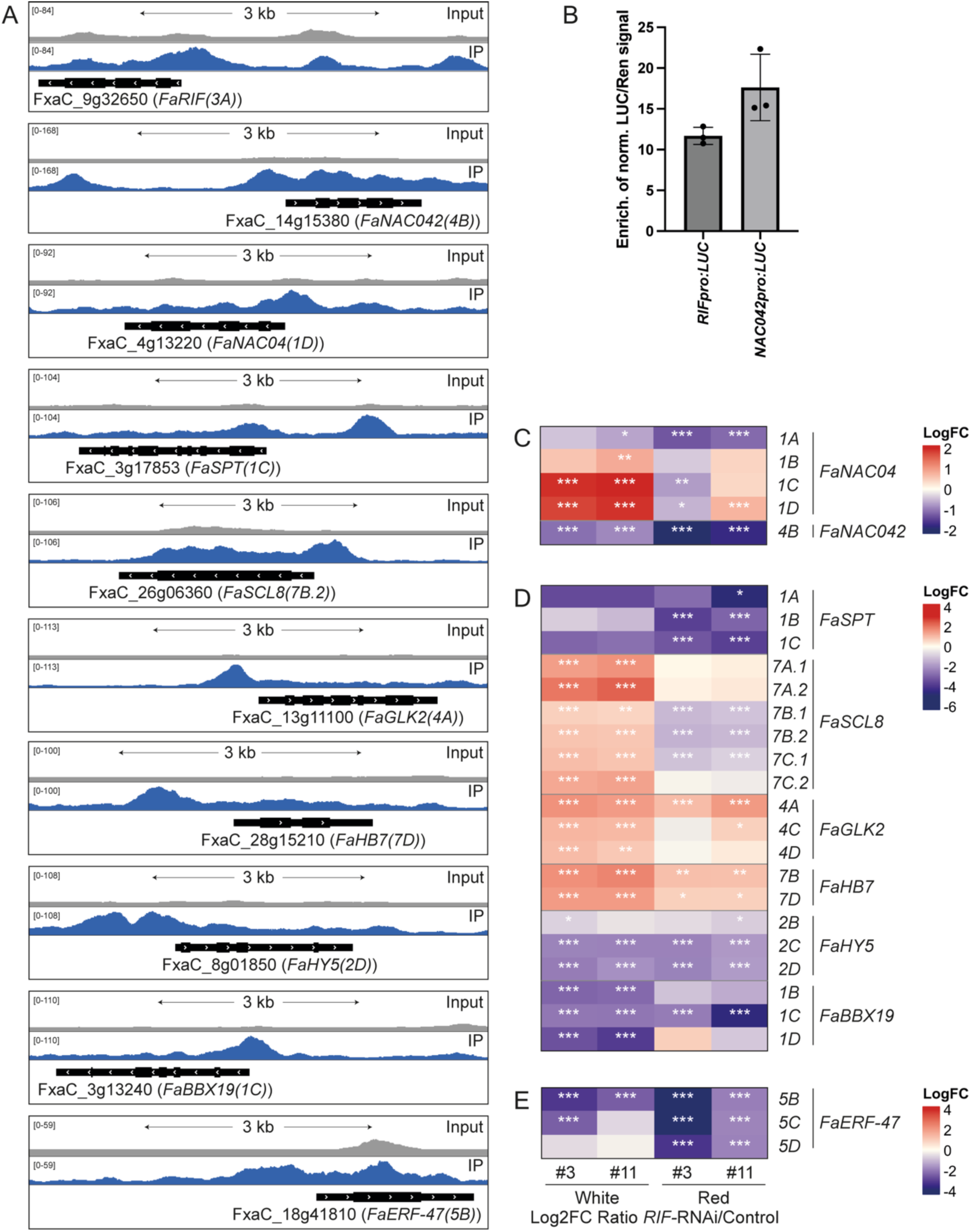
Transcription factors directly regulated by FaRIF. **A)** ChIP-seq peaks showing FaRIF binding sites and input reads over representative homoeologs of ripening-related TFs. **B)** *FaRIF* and *FaNAC042* promoters activation by dual-luciferase reporter assay in infiltrated *N. benthamiana* leaves. Firefly LUCIFERASE (LUC) activity is presented relative to RENILLA (REN) activity and normalized against the control (empty effector vector). Each value represents a biological replicate, each with 3 technical replicates. **C) to E)** Heatmaps displaying the expression of FaRIF-regulated transcription factors. Panels include transcription factors from the NAC family (C), involved in fruit development and ripening-related processes (D), and aroma compounds biosynthesis (E). Significant DEGs are marked with asterisks (***FDR≤0,0005; **FDR≤0,005; *FDR≤0,05). Detailed information is available in Supplementary Data Set S7.

Other TFs previously reported to regulate strawberry fruit development and ripening were identified as FaRIF direct targets (Fig. 4D, Supplementary Fig. S5B and Data Set S7). These included all homoeologs of the bHLH TF *FaSPT*, involved in strawberry fruit development (Tisza et al., 2010) and sugar accumulation during ripening (Chen et al., 2024), with three homoeologs downregulated in the *FaRIF*-RNAi lines. Similarly, all homoeologs of *FaSCL8*, encoding a GRAS protein that regulates anthocyanin, soluble sugar accumulation, and fruit firmness (Pillet et al., 2015; Zheng et al., 2023) were identified as direct targets, with some downregulated at the red stage. All homoeologs of the MYB TF *GOLDEN2-LIKE* (*FaGLK2*), also known as *FvMYBR139* in *F. vesca* (Li et al., 2021), were also identified as direct targets and upregulated in the RNAi lines, suggesting that FaRIF may directly and positively regulate strawberry degreening during ripening, as the tomato ortholog of this TF promotes plastid formation and high chlorophyll levels (Nguyen et al., 2014). Additionally, all homoeologs of the homeodomain-leucine zipper (HD-Zip) class I TF *FaHB7*, whose Arabidopsis ortholog, *AtHB7*, acts as a negative regulator of ABA signaling (Valdés et al., 2012), were significantly upregulated at both stages, with two identified as direct targets. Furthermore, most homoeologs of the bZIP *FaHY5* and the B-box gene *FaBBX19*, both associated with ABA signaling in Arabidopsis (Chen et al., 2008; Bai et al., 2019), were downregulated at both stages, with three identified as direct FaRIF targets. These findings further support FaRIF’s role in modulating ABA signaling, contributing to the delayed ripening phenotype observed in the *FaRIF-*RNAi lines (Fig. 1C) (Martín-Pizarro et al., 2021).

In addition to the direct targets, several important TFs involved in the biosynthesis of anthocyanins and/or proanthocyanidins (PA) were found indirectly regulated by FaRIF (Supplementary Fig. S5C, Supplementary Data Set S7). Among them, we identified homoeologs of the MADS-box TF *FaSHP* (Daminato et al., 2013) and the MYB TFs *FaMYB52*, whose Arabidopsis ortholog, *AtMYB4*, regulates anthocyanin and proanthocyanidin (PA) biosynthesis (Wang et al., 2020), *FaMYB77* (Xu et al., 2021), *FaMYB10(1B)*, the dominant homoeolog of the key regulator of anthocyanin biosynthesis (Castillejo et al., 2020), and *FaMYB123* (*F.* vesca’s *FvMYB79* (Cai et al., 2021)) (Martínez-Rivas et al., 2023; Cai et al., 2021). Additionally, homoeologs of the B-box gene *FaBBX22*, known to cooperate with *FaHY5* in light-induced anthocyanin biosynthesis (Liu et al., 2022b), as well as *FaPRE1*, an atypical HLH TF promoting anthocyanin biosynthesis in leaves when stably overexpressed (Medina-Puche et al., 2021), were found differentially expressed. Together, these results support that the altered flavonoid, and particularly anthocyanin, content in the *FaRIF*-RNAi lines might be explained by the broad disturbance in the expression of all these regulators.

Furthermore, TFs involved in volatile organic compounds (VOCs) biosynthesis were also found among the direct and indirect targets of FaRIF (Fig. 4E, Supplementary Fig. S5D and Data Set S7). Thus, homoeologs of *FaDOF2* and *FaEOBII*, encoding DOF-type and MYB TFs, respectively, which synergistically enhance eugenol biosynthesis (Medina-Puche et al., 2015; Molina-Hidalgo et al., 2017), were indirectly downregulated at the red stage. Furthermore, homoeologs of *FaERF-47*, an AP2/ERF TF linked to volatile esters biosynthesis through activation of the acyltransferase-encoding gene *AAT* in *F. vesca* (Li et al., 2020), were identified as direct targets of FaRIF and significantly downregulated at both ripening stages.

In summary, these results highlight a complex regulation of the strawberry fruit development and ripening processes, with FaRIF emerging as a more prominent upstream direct and indirect regulator of several important ripening-related TFs than previously reported in our earlier work (Martín-Pizarro et al., 2021).

### FaRIF directly regulates cell wall remodeling genes

Within the “Cell wall organization” functional category enriched among the DEGs and FaRIF-direct targets, we identified numerous genes encoding Expansins (EXP), proteins involved in cell wall loosening (Cosgrove, 1998; Dotto et al., 2006). Eight *EXP* genes were mainly downregulated at either white or red stages, or both, depending on the gene. Notably, all homoeologs of *FaEXP1* and some of the most highly expressed *EXPs* during ripening in *F. vesca* (Dong et al., 2022) and *F.* × *ananassa*, as *FaEXP2(7D)*, the most expressed *EXP* gene, were identified as direct targets of FaRIF (Fig. 5A, D, Supplementary Fig. S6 and Data Set S7), supporting a direct role for this TF in promoting cell wall modification to favor strawberry fruit softening.

**Figure 5.**
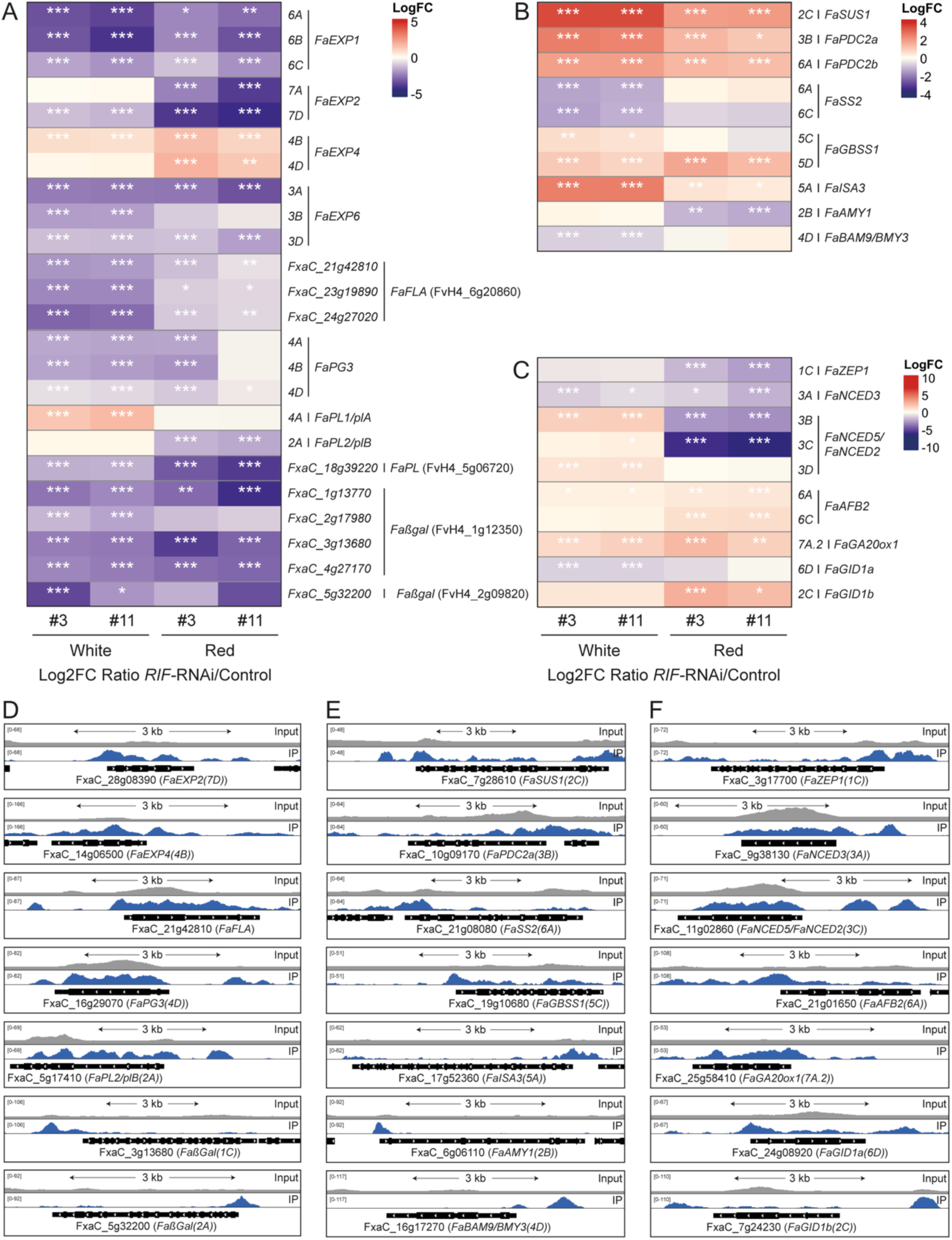
FaRIF directly regulates genes involved in cell wall modification, carbohydrate metabolism, and hormone biosynthesis and signaling pathways. **A) to C)** Heatmaps displaying the expression of FaRIF direct regulated genes related to cell wall modification **(A)**, carbohydrate metabolism **(B)**, and hormone biosynthesis and signaling pathways (C). Significant DEGs are marked with asterisks (***FDR≤0,0005; **FDR≤0,005; *FDR≤0,05). **D) to F)** ChIP-seq peaks showing FaRIF binding sites and input reads over representative homoeologs of direct targets in cell wall modification **(D)**, carbohydrate metabolism **(E)**, and hormone-related genes **(F)**. Detailed information is available in Supplementary Data Set S7.

In addition to Expansins, nine genes encoding Fasciclin-like arabinogalactan proteins (FLAs), involved in cell adhesion, were significantly downregulated in both silenced lines, with homoeologs of *FaFLA17* (FvH4_6g20860 in *F. vesca*) identified as direct targets of FaRIF (Fig. 5A, D, Supplementary Fig. S6 and Data Set 7). Genes encoding pectin-modifying enzymes were also primarily downregulated. Among these were genes encoding nine pectin methylesterases (*FaPMEs*), including *FaPME39*, known to promote fruit softening in *F. vesca* (Xue et al., 2020) (Supplementary Fig. S6 and Data Set 7). Furthermore, most homoeologs of the polygalacturonase-encoding gene *FaPG3*, as well as homoeologs of the pectate lyases *FaPL1/plA* and *FaPL2/plB*, both known to promote strawberry fruit softening (Medina-Escobar et al., 1997; Jiménez-Bermúdez et al., 2002; Benítez-Burraco et al., 2003; Zhang et al., 2022b), along with two additional *FaPL* genes were identified as direct targets of FaRIF (Fig. 5A, D, Supplementary Fig. S6 and Data Set 7).

Lastly, homoeologs of genes encoding rhamnogalacturonan-I-modifying enzymes, in particular of four β-galactosidase genes, including *Fa*β*Gal1* and *Fa*β*Gal3* (Trainotti et al., 2001), were also identified as direct targets of FaRIF (Supplementary Data Set 7). Notably, all homoeologs of another *FaGal* gene (FvH4_1g12350 in *F. vesca*) contained FaRIF binding sites, while two other *FaGal* genes, including *Fa*β*Gal4*, which promotes strawberry fruit softening (Paniagua et al., 2016), were indirectly regulated by FaRIF (Fig. 5A, D, Supplementary Fig. S6 and Data Set 7). Altogether, these data highlight FaRIF’s role in regulating cell wall-related genes critical for promoting fruit softening during strawberry fruit ripening, consistent with the increased fruit firmness observed in *RIF*-silenced and knockout lines in *F.* × *ananassa* and *F. vesca*, respectively (Martín-Pizarro et al., 2021; Li et al., 2023).

### FaRIF regulates carbohydrate metabolism

Within the “Carbohydrate metabolism” category, homoeologs of genes encoding enzymes involved in local sucrose metabolism were identified among the FaRIF regulated genes (Fig. 5B, E, Supplementary Fig. S7A and Data Set S7) (Luo et al., 2020). Several *SUCROSE SYNTHASE1* (*FaSUS1*) homoeologs, which degrade sucrose into UDP-glucose and fructose, were upregulated at both ripening stages, with one containing a FaRIF binding site. In contrast, the only expressed *SUCROSE PHOSPHATE SYNTHASE1* (*FaSPS1*) homoeolog, *FaSPS1(2B)*, involved in sucrose biosynthesis, was downregulated in both ripening stages. This aligns with the altered sugar content in the *FaRIF*-RNAi receptacles, where glucose and fructose levels increased, while sucrose content is reduced in ripe *FaRIF-*RNAi receptacles (Martín-Pizarro et al., 2021).

Fruit starch metabolism is also important for the final sugar content in ripe strawberries (Souleyre et al., 2004), being degraded throughout the ripening process by amylases. Interestingly, homoeologs of *STARCH SYNTHASE 2* (*FaSS2*) were identified as direct targets of FaRIF, being downregulated at the white stage in the *FaRIF*-silenced lines. Additionally, most homoeologs of *GRANULE BOUND STARCH SYNTHASE 1* (*FaGBSS1*), involved in amylose and starch biosynthesis in peas and Arabidopsis (Denyer et al., 1999; Seung et al., 2015), were also identified as direct targets of FaRIF, being in this case mainly upregulated in the silenced lines. Furthermore, genes encoding for α- and β-amylases, as well as for an isoamylase, *FaISA3*, were also found to be misregulated when *FaRIF* was silenced, with some of them being among FaRIF’s direct targets. Thus, these results suggest that the FaRIF not only controls sugar content during strawberry ripening by regulating sucrose metabolism, as previously reported (Martín-Pizarro et al., 2021; Li et al., 2023) but also plays a role in controlling starch metabolism, contributing to the final sugar content of the ripe strawberry fruits.

MapMan analysis of the DEGs confirmed the enrichment in genes involved in the fermentation process at both ripening stages. Notably, two PYRUVATE DECARBOXYLASE-encoding genes (*FaPDC*s) were upregulated at both ripening stages in the *FaRIF-*silenced fruits, with one homoeolog of each identified as a direct target of FaRIF. Indirect regulation was also observed for most homoeologs of two *ALCOHOL DEHYDROGENASE* genes (*FaADHs*), which were mainly upregulated at both ripening stages. These findings support a direct role of FaRIF in regulating the aerobic/anaerobic balance, which is known to change during strawberry fruit ripening (Wang et al., 2017).

### FaRIF regulates phytohormone biosynthesis and signaling

Within the “Phytohormone action” MapMan category, the ABA-related subcategory was enriched in *FaRIF*-RNAi fruits at the ripe stage and among the FaRIF direct targets (Fig. 3D). Thus, genes involved in the rate-limiting steps of ABA biosynthetic pathway, such as *zeaxanthin epoxidase 1* (*FaZEP1*), and *9-cis-epoxycarotenoid dioxygenases* (*FaNCEDs*), were identified as FaRIF-direct targets and, along with *neoxanthin synthase* (*FaNSY*), were significantly downregulated in the *FaRIF-*silenced fruits (Fig. 5C, F, Supplementary Fig. S7B and Data Set 7). Furthermore, homoeologs of ABA receptors such as *FaPYL2* (PYRL/PYL) and *FaABAR* (ABA receptor/Mg-chelatase H subunit (ABAR/CHLH)), both known to positively influence strawberry fruit ripening (Jia et al., 2011; Hou et al., 2018; 2021; Li and Shen, 2023), were positively regulated by FaRIF, being one of the *FaABAR* homoeologs a direct target (Supplementary Fig. S7B and Data Set S7). Additionally, protein kinase-encoding genes *SUCROSE NONFERMENTING1-RELATED PROTEIN KINASE 1*, (*FaSnRK1s*), involved in ABA signaling (Jossier et al., 2009), were found enriched among the FaRIF targets and differentially expressed, mainly at the white stage (Fig. 3D, Supplementary Fig. S7B and Data Set S7). All these findings are consistent with the altered ABA content and delayed ripening phenotype of *FaRIF-*silenced fruits (Martín-Pizarro et al., 2021), supporting the positive role of FaRIF in regulating the key phytohormone involved in strawberry fruit ripening (Jia et al., 2011).

Auxins and gibberellins are crucial for promoting strawberry fruit growth and inhibiting ripening (Perotti et al., 2023) by suppressing ABA accumulation through *CYP707A4a* expression activation (Liao et al., 2018). Interestingly, genes related to the content and signaling pathways of these phytohormones were also misregulated in the *FaRIF*-silenced fruits (Fig. 5C, F, Supplementary Fig. S7B and Data Set S7). Direct targets of FaRIF, such as most homoeologs of the auxin receptor gene *AUXIN SIGNALING F-BOX 2* (*FaAFB2*), which positively mediates auxin signaling in Arabidopsis (Lakehal et al., 2019), were upregulated at the red stage, while the negative regulator *FaAUX/IAA13* was repressed at the white stage, suggesting a negative role of FaRIF in regulating auxin signaling. FaRIF may also repress the expression of the GA biosynthetic gene *FaGA20ox1*, while *FaRIF*-silencing led to opposing indirect regulation of two homoeologs of the GA catabolic gene *FaGA2ox1* (Hytönen et al., 2009; Csukasi et al., 2011). Additionally, homoeologs of two *GIBBERELLIN-INSENSITIVE DWARF1* (*FaGID1a* and *FaGID1b*) (Csukasi et al., 2011), involved in GA signaling, were identified as FaRIF-direct targets with opposing regulation. These results suggest that, in addition to regulating ABA, FaRIF might contribute to strawberry fruit ripening controlling also auxin and GA biosynthesis and signaling.

### FaRIF regulates genes involved in secondary metabolism

The “secondary metabolism” bin was also enriched at the red stage in the *FaRIF-* silenced lines. Interestingly, genes involved in terpenoid biosynthesis were identified, including homoeologs involved in the first committed steps in the mevalonate (MVA) and the methylerythritol phosphate (MEP) pathways, such as 3-hydroxy-3-methylglutaryl-CoA reductases (HMGRs) and 1-deoxy-D-xylulose 5-phosphate synthase (DXS), respectively. These genes were indirectly misregulated in a manner where those normally induced during ripening were downregulated, while those usually repressed during this process were upregulated (Supplementary Figure S8 and Data Set S7). Additionally, two homoeologs of a terpene synthase, *FaTPS11* (Zhang et al., 2022a), were identified as FaRIF-direct targets and downregulated at the ripe stage in the *FaRIF-*silenced fruits. Moreover, the four homoeologs of the *PHYTOENE SYNTHASE 1* (*FaPSY*), which initiates the carotenoid biosynthesis, were also downregulated at the red stage, with two being FaRIF direct targets. This suggests a novel role of FaRIF in regulating terpenic compounds biosynthesis.

A specific bin related to phenolic compounds was also enriched at the red stage, although only a few were identified as FaRIF direct targets (Fig. 6, Supplementary Figure S9 and Data Set S7). Indirect regulation was observed for most structural genes in the phenylpropanoid and flavonoid pathways, including those encoding phenylalanine ammonia-lyases 1 and 2 (*FaPAL1* and *FaPAL2*), chalcone synthase 1 (*FaCHS1*), chalcone isomerase 2 (*FaCHI2*), flavanone 3-hydroxylase (*FaF3H*), dihydroflavonol 4-reductase (*FaDFR*), anthocyanidin synthase (*FaANS/LDOX*), and glutathione S-transferase (GST), as well as Reduced Anthocyanins in Petioles-Like 1 (*FaRAP-L1*), an anthocyanin transporter to the vacuole that promotes strawberry fruit pigmentation (Luo et al., 2018). On the contrary, all homoeologs of *Fa4CL1*, one of the two paralogs encoding for 4-Coumaroyl-CoA ligases found downregulated in the *FaRIF-*RNAi lines, were identified as direct FaRIF targets.

**Figure 6.**
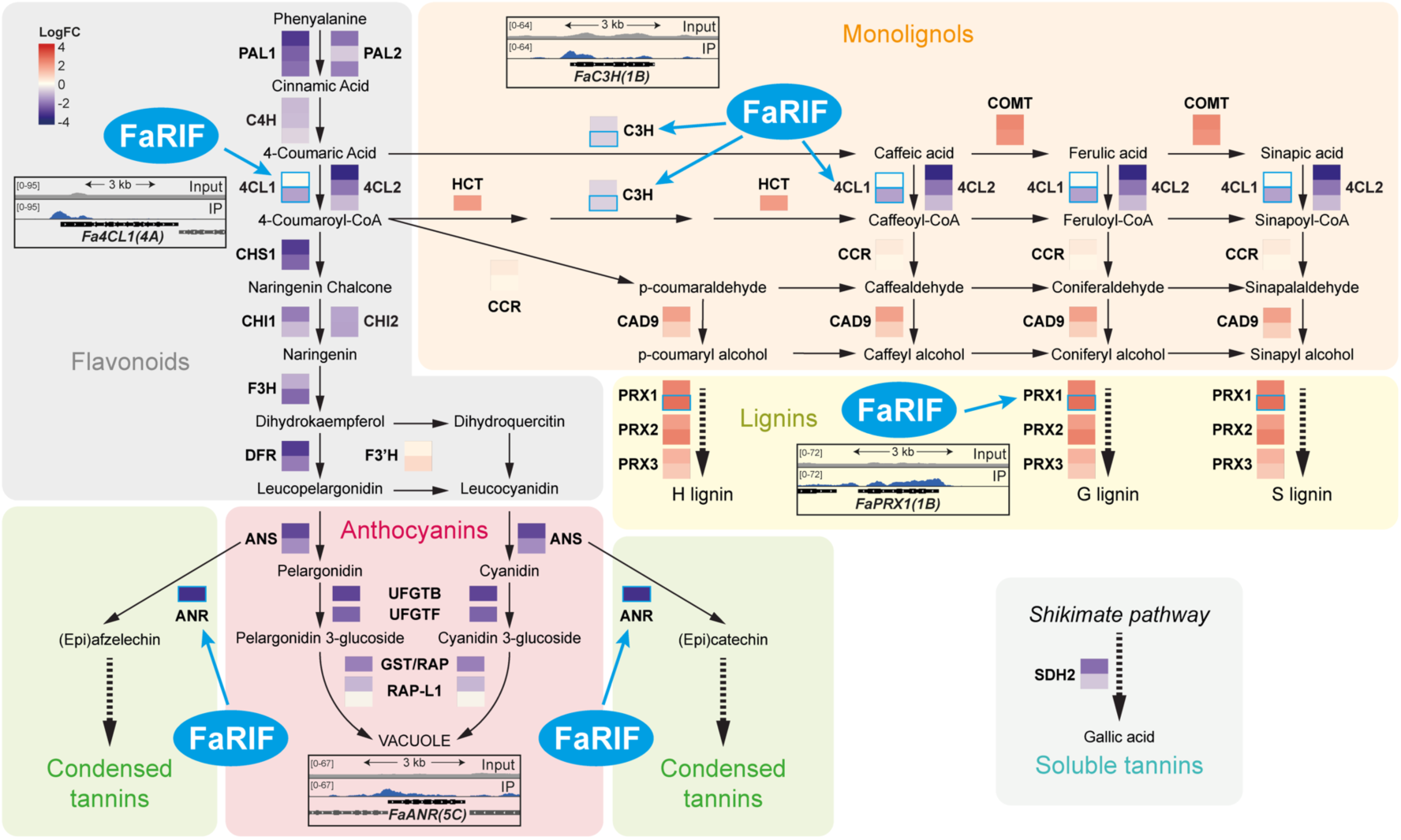
FaRIF indirectly and directly regulates genes involved in the phenylpropanoid, flavonoid, monolignol, lignin, condensed and soluble tannins pathways. Colors denote the average log2 expression ratio of *35Spro:RIF*-RNAi/control for differentially expressed homoeologs of each gene (bold fonts) in both transgenic lines at the red stage. Red and purple show up- and downregulation, respectively, in both silenced lines. Direct targets are depicted with a blue frame, and their binding sites shown in the ChIP-seq peaks insets. Detailed information is available in Supplementary Data Set S7.

Similarly, other structural genes in the phenylpropanoid, flavonoid, and monolignol pathways were also misregulated in the *FaRIF*-silenced fruits, with some homoeologs of 4-Coumarate 3-hydroxylase (*FaC3H*) and a peroxidase (*FaPRX1*) being direct targets of FaRIF. These results align with previous observations supporting a positive role of FaRIF regulating anthocyanin content (Fig. 1C) (Martín-Pizarro et al., 2021; Li et al., 2023) and a negative role in lignification (Martín-Pizarro et al., 2021), contributing to strawberry fruit color and softening during ripening.

### FaRIF forms homo- and heterodimers with NAC TFs and other proteins

Our transcriptomic and ChIP-seq data highlighted the central role of FaRIF in multiple processes essential for fruit ripening. NAC proteins are known to modulate their activity by forming homo- or heterodimers (Xie et al., 2000; Hegedus et al., 2003; Ernst et al., 2004; Ren et al., 2021; DIncà et al., 2023; Lu et al., 2024a), and by interacting with other proteins, such as bZIP-type (Xu et al., 2013) and homeodomain (HD) TFs (Liu et al., 2019), as well as enzymes like phosphatases (Guan et al., 2014) and kinases (Li et al., 2023). These findings prompted us to investigate the potential involvement of different protein partners in mediating FaRIF’s diverse biological functions. To this end, we employed a TurboID-based proximity labeling approach to screen for interacting proteins of FaRIF, using the Ti-TAN plasmid toolbox (Tan et al., 2024). First, the nuclear localization of the *FaRIF-TurboID-GFP* fusion protein was confirmed by transient expression in *N. benthamiana* leaves (Supplementary Fig. S10A). Next, the final *35Spro:FaRIF-3xHA-TbID* vector was generated and the efficiency of the system in strawberry fruits was tested by transient expression in fruits at the initial white stage (Supplementary Fig. S10D). Once the fruits ripened to the red stage, they were infiltrated with 50 µM of biotin solution, and biotinylation was confirmed by western blot (Supplementary Fig. S10B). Interestingly, ripe strawberry fruits naturally contain high biotin levels (around 15 ng/g fresh weight (Staggs et al., 2004)), a characteristic that distinguishes them from commonly used model species for proximity labeling assays, such as *N. benthamiana* and *A. thaliana*. Thus, despite biotin supplementation, no significant differences in global protein biotinylation were observed between treated and untreated fruits.

To identify proteome-wide interactors of FaRIF, we generated three independent biological replicates per construct (*35Spro:FaRIF-3xHA-TbID* and *35Spro:3xHA-TbID-3xNLS* as a negative control) (Supplementary Figs. S10C and S10D). Candidate interacting proteins identified by LC-MS/MS from the *FaRIF-TbID* samples were compared to those from the negative *TbID-3xNLS* control to eliminate those overlapping, and therefore, non-specific biotinylated proteins. We confidently identified a total of 973 peptides corresponding to 43 protein groups, representing 247 unique proteins that were exclusively present in the *FaRIF-TbID* samples across the three biological replicates (Supplementary Data Set S8). Interestingly, several NAC TFs close to FaRIF, which identified peptides are common for all of them, were found among the potential interactors, including FaRIF itself as well as FaNAC004, FaNAC015, FaNAC021, FaNAC034, FaNAC039, FaNAC040, FaNAC045, FaNAC064, and FaNAC077 (Supplementary Data Set S8), suggesting that FaRIF may form both homo- and heterodimers with other close NAC family members.

In addition, other putative interactors included several ribosomal and RNA-binding proteins. Among these was the putative ortholog of Arabidopsis PUF RNA-binding protein PUMILIO2 (PUM2), involved in translational regulation, mRNA localization (Quenault et al., 2011) and pre-ribosomal RNA processing (Abbasi et al., 2010; Qiu et al., 2014), as well as CTC-Interacting Domain 4 (CID4), which is involved in developmental pathways and stabilizes target mRNAs via 3’UTR binding (López-Juárez et al., 2021). Several YTH domain-containing proteins, including FaYTH3, FaYTH4, FaYTH6, and FaYTH8, orthologs of *F. vesca* proteins recently identified (Xu et al., 2023) and reported to regulate mRNA splicing, processing, stability, and translation (Flores-Téllez et al., 2023), were also identified.

Proteins related to protein folding and processing also emerged as FaRIF interactors, notably CCT3, a component of the chaperonin TCP-1 ring complex (TRiC/CCT) essential for the proper folding of specific eukaryotic proteins (Spiess et al., 2004), and UBIQUITIN-SPECIFIC PROTEASE 7 (UBP7), which cleaves ubiquitin chains to likely prevent substrate degradation (Zhou et al., 2017; Yan et al., 2000), suggesting a role for these proteins in FaRIF protein homeostasis. Finally, we identified two Ca^2+^-dependent lipid-binding (CalB) proteins, which generally bind calcium and phospholipids (Maguire et al., 2024), specifically FaCalB1 and FaCalB2, each containing a single C2 domain.

We next selected several of these interactors identified by proximity labeling for further validation, focusing on FaRIF itself, as well as FaNAC021 and FaNAC034. Both NAC proteins are encoded by homoeologs upregulated throughout ripening, suggesting their potential involvement in this process (Fig. 7A). Additionally, we included C2 domain protein FaCalB1, whose expressed homoeologs all correlate with *FaRIF* expression, and the chaperonin FaCCT3 (Fig. 7A). Transient expression of RFP-tagged versions of these putative interactors in *N. benthamiana* leaves confirmed the colocalization with FaRIF-GFP in nuclei, except for FaCalB1, which also exhibited cytoplasmic localization, and FaCCT3, which was predominantly localized in the cytoplasm with limited nuclear presence (Fig. 7B). Co-immunoprecipitation (CoIP) assays subsequently confirmed the interactions between FaRIF and all selected candidates (Fig. 7C). Notably, FaRIF homo- and heterodimerization with FaNAC021 and FaNAC034 showed the strongest interactions, while weaker interactions were detected with FaCalB1 and FaCCT3, despite FaCCT3’s low nuclear signal (Fig. 7C). These results confirm that these proteins identified via proximity labeling interact with FaRIF and may contribute to the regulation of its activity and/or protein homeostasis.

**Figure 7.**
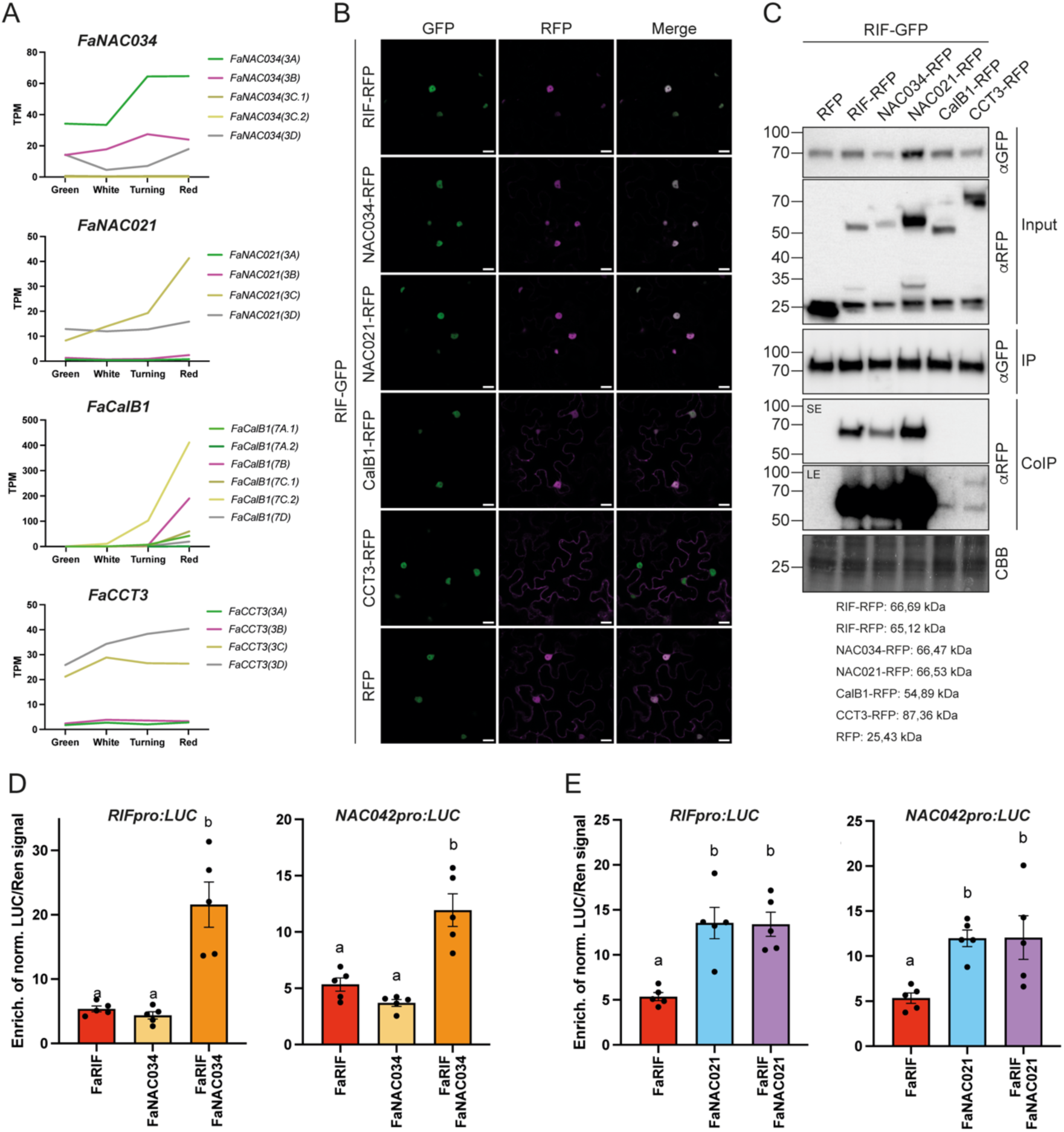
Validation of FaRIF interactions with other proteins. **A)** Expression patterns of the homoeologs of candidate FaRIF interactors in *F.* × *ananassa* receptacles at four ripening stages. Data adapted from (Sánchez-Sevilla et al., 2017; Liu et al., 2021). **B)** Subcellular localization of FaRIF-GFP and RFP-tagged candidate interactors in *N. benthamiana* leaves. Images were captured 48h post-infiltration. Scale bars = 20 µm. **C)** Co-immunoprecipitation (CoIP) assay of FaRIF-GFP with RFP-tagged FaRIF, FaNAC034, FaNAC021, FaCalB1, and FaCCT3. All tagged proteins were transiently coexpressed in *N. benthamiana* leaves, and FaRIF-GFP was immunoprecipitated using anti-GFP-Trap beads. Total (input), IP, and CoIP proteins were analyzed by immunoblotting. Equal loading was confirmed by Coomassie Brilliant Blue (CBB) staining of input samples. GFP- and RFP-tagged proteins were detected using anti-GFP (αGFP) and anti-RFP (αRFP) antibodies, respectively. SE and LE denote shorter and longer exposure times, respectively. **D, E)** Dual-luciferase reporter assays testing the activation of the *RIF* and *NAC042* promoters in *N. benthamiana* leaves. The individual activity of FaRIF and its combined activity with FaNAC034 **(D)** and FaNAC021 **(E)** were tested. Firefly LUCIFERASE (LUC) activity is presented relative to RENILLA (REN) activity and normalized against the control (empty effector vector). Each value represents the mean of 3 technical replicates for 5 biological replicates ±SD. Different letters indicate significant differences at P≤0,05.

### FaRIF activity is modulated by FaNAC034

Our previous data confirmed that FaRIF can form both homo- and heterodimers, at least with FaNAC034 and FaNAC021. Thus, we next investigated whether the formation of these heterodimers influences FaRIF’s transcriptional activity on specific target genes. To test this, we selected two validated FaRIF-bound promoters (its own promoter and that of *FaNAC042*) and confirmed that FaNAC034 and FaNAC021 also activated them in a dual-luciferase transactivation assay in *N. benthamiana* leaves (Fig. 7D, E). Interestingly, while FaNAC034 activated both promoters similarly to FaRIF (Fig. 7D), FaNAC021 induced significantly higher expression on those promoters than FaRIF (Fig. 7E). Then, to explore the effects of heterodimerization on the activation of these promoters, we coinfiltrated FaRIF with either FaNAC034 or FaNAC021 in the transactivation assay. Notably, the FaRIF-FaNAC034 combination led to greater activation of both promoters compared to the individual NACs alone, suggesting a synergistic effect of this heterodimer. In contrast, the FaRIF-FaNAC021 combination did not enhance activation beyond that of FaNAC021 alone, indicating that, at least for these two promoters, FaRIF does not increase the activity of FaNAC021 when forming a heterodimer. Further experiments are needed to better understand the broader impact of FaRIF heterodimerization on its transcriptional activity and to assess the extent of the overlap among the target genes of these three NAC TFs genome-wide.

## Discussion

In this work, we have established optimized protocols to identify genome-wide binding sites and the proxitome of FaRIF, a key TF in strawberry ripening (Martín-Pizarro et al., 2021; Li et al., 2023). These approaches allowed us to elucidate the gene regulatory network directly and indirectly modulated by FaRIF, and a set of protein interactors influencing its transcriptional activity.

### Insights from using the octoploid genome as a reference for transcriptome analysis in *F.* × *ananassa*

*RIF* expression increases throughout ripening in both *F.* × *ananassa* and *F. vesca* receptacles (Martín-Pizarro et al., 2021; Li et al., 2023). However, RNA-seq data from receptacles at four ripening stages mapped to the *F.* × *ananassa* reference genome (Liu et al., 2021) revealed that *FaRIF(3D)*, encoded by the *F. viridis* subgenome, did not follow this trend, while *FaRIF(3A)*, from the *F. vesc*a subgenome, was the most expressed homoeolog (Fig. 1B). ChIP-seq data revealed that FaRIF’s preferential binding to the *F. vesca* subgenome (Fig. 2E, Supplementary Fig. S3), consistent with its dominance in *F.* × *ananassa* due to lower transposable elements density and higher homoeologs expression (Edger et al., 2019). Nevertheless, exceptions exist. For instance, the most expressed homoeolog of *FaEXP1*, *FaPME39, FaNCED5/FaNCED2*, and *FaCHS1* is encoded by another subgenome, following also their respective homoeologs different expression trends (Supplementary Data Set S7). Thus, although the extended use of the diploid *F. vesca* genome as the reference for transcriptome analysis in the octoploid species has proven successful and informative, the availability of a high-quality reference genome for *F.* × *ananassa* marks a significant advancement. Despite the analytical challenges posed by the complexity of the octoploid genome, including the presence of homoeologs for each gene, the use of *F.* × *ananassa* genome provides a more accurate and comprehensive view of the transcriptomic landscape in this economically highly valuable species, being indispensable for achieving a bona fide understanding of gene expression and regulation in *F.* × *ananassa*.

### *FaRIF* directly regulates key ripening-related processes

Combined DEG and ChIP-seq analyses revealed that FaRIF directly or indirectly regulates genes involved in key ripening processes, including ABA biosynthesis and signaling, sugar metabolism, secondary metabolites biosynthesis, cell wall remodeling, and energy balance. Among the DEGs in *FaRIF-*RNAi lines, 15.3% were identified as direct targets, consistent with findings for its *F. vesca* ortholog (Fig. 3C, Supplementary Fig. S4A) (Li et al., 2023). These results underline FaRIF’s central role as an upstream regulator of strawberry ripening by influencing a wide array of processes, either directly or via the regulation of other TFs, such as FaSPT and FaSCL8, which are known regulators of ripening. Notably, FaRIF directly regulates genes involved in ABA biosynthesis and signaling, the primary hormone modulating strawberry fruit ripening (Chai et al., 2011; Jia et al., 2011; Li et al., 2022a). Notably, the ABA signaling-related SnRK1-encoding genes, which have also been associated with sugar accumulation in strawberry (Luo et al., 2020) and ABA-mediated seed maturation in pea (Radchuk et al., 2006), were enriched among the direct targets of FaRIF. This supports the critical role of FaRIF orchestrating ripening, as evidenced by the complete ripening block in *Fvrif* knockout *F. vesca* mutants (Li et al., 2023).

Our results show that FaRIF’s influence on secondary metabolism, including flavonoids and lignin biosynthesis may be mediated through direct regulation of *Fa4CL1*, *FaANR*, *FaC3H*, and *FaPRX1*. While previous DAP-seq analysis identified *FvMYB10* as a direct target of FvRIF in *F. vesca* (Li et al., 2023), this regulation was not observed in *F.* × *ananassa*. This discrepancy may reflect divergent roles of RIF orthologs between species, where the regulation of *MYB10* in *F. vesca* has a stronger impact on anthocyanin biosynthesis than in *F.* × *ananassa.* Alternatively, it could be due to a false-negative result in our ChIP-seq analysis. Furthermore, our data revealed a role of FaRIF in the regulation of terpenoid biosynthesis, including VOCs and carotenoids. Among the direct targets, *FaTPS11* is the closest paralog to *NEROLIDOL SYNTHASE1* (*FaNES1*), the monoterpene linalool and the sesquiterpene nerolidol (Aharoni et al., 2004). These VOCs contribute with sweet and flowery notes to strawberry aroma (Ulrich et al., 1997). While *FaTPS11* has not yet been functionally characterized, its homoeologs were identified as candidate genes for the biosynthesis of these VOCs in a multiomic QTL analysis (Barbey et al., 2020). Thus, we hypothesize that FaRIF directly promotes *FaTPS11* expression, contributing to linalool and nerolidol biosynthesis and enhancing aroma-related traits.

Remarkably, our ChIP-seq study has allowed the identification of new FaRIF targets not previously reported in the DAP-seq analysis in *F. vesca* (Supplementary Table S2), including FaRIF itself, which suggests a feedback positive regulatory loop. We also identified novel FaRIF targets involved in ABA biosynthesis and signaling, (e,g., *FaZEP1*, *FaNCED3*, *FaHY5*, *FaBBX1*, *FaHVA22*, and *FaSnRK2.6*), cell wall modification (e.g., *FaEXP1*, *FaPG3*, *FaPL3*, and *FaPL4*), and in quality traits related to sugar, ascorbic acid, and eugenol biosynthesis (e.g., *FaSUS1*, *FaVTC2*, and *FaDOF2*).

### *FaRIF* interacts with ripening-related NAC TFs and proteins involved in mRNA and protein homeostasis

NAC proteins are known not only to homo- and heterodimerize (Xie et al., 2000; Hegedus et al., 2003; Ernst et al., 2004; Zhou et al., 2015; Gladman et al., 2016; Mathew et al., 2016; Ren et al., 2021; DIncà et al., 2023; Lu et al., 2024a), but also to interact with other proteins, such as bZIP-type and homeodomain (HD) TFs (Xu et al., 2013; Liu et al., 2019), and enzymes like phosphatases and kinases (Guan et al., 2014; Li et al., 2023) that regulate their activity. Using TurboID, we found that FaRIF interacts with other NAC TFs, including FaRIF itself, FaNAC021 and FaNAC034. CoIP validation confirmed the formation of both homo- and heterodimers. Notably, FaRIF-FaNAC034 heterodimers showed a synergistic effect in activating the gene promoters tested. Similar synergistic effects among NAC TFs have been previously reported, such as in grapevine, where combinations of VviNAC03, VviNAC33 and VviNAC60 enhance transcription of genes involved in chlorophyll degradation and anthocyanin accumulation (DIncà et al., 2023), and in *N. tabacum*, where NtNAC028 and NtNAC080 synergistically promote jasmonic acid biosynthesis by regulating *NtLOX3* expression (Lu et al., 2024a). Further studies are required to determine the extent of genes coregulated by FaRIF, FaNAC021, and FaNAC034, and how FaRIF’s interactions with these NAC TFs fine-tune target gene expression during ripening.

In addition to NAC TFs, our TurboID-based proximity labeling analysis revealed ribosomal proteins as potential interactors. While these may reflect biotinylation during translation of the FaRIF-TbID protein, functional complexes with these proteins cannot be ruled out. Interestingly, FaRIF was found to interact with RNA-binding proteins, including PUM2, which is involved in translational regulation, mRNA localization, and pre-ribosomal RNA processing in Arabidopsis (Quenault et al., 2011; Abbasi et al., 2010; Qiu et al., 2014). PUM proteins have not been studied in fruit ripening, but related proteins such as APUM5 and APUM9 modulate ABA-responsive genes (Huh and Paek, 2014; Nyikó et al., 2019). This raises the possibility that FaRIF-PUM interactions contribute to ABA signaling regulation in strawberry ripening, a hypothesis for further exploration.

Other identified RNA-binding proteins included YTH domain-containing proteins, which recognize N6-methyladenosine (m^6^A) modifications in mRNAs and influence their stability and translation (Zhou et al., 2021; Flores-Téllez et al., 2023). In strawberry, changes in m^6^A methylation occur at the onset of strawberry ripening, stabilizing ABA-related mRNAs that encode the rate-limiting enzyme *NCED5* and the transcription factor *AREB1*, while also facilitating the translation of *ABAR*, an ABA receptor (Zhou et al., 2021). SlYTH1 and SlYTH2, orthologs in tomato of the FaRIF interactors FaYTH8 and FaYTH3, respectively, have been implicated in the development and ripening of the fruit (Yin et al., 2022; Ao et al., 2023). Notably, *SlYTH2* knockout mutants display delayed internal fruit ripening and altered ABA content (Ao et al., 2023). Furthermore, two *FaYTH4* homoeologs coexpressed with FaRIF (Supplementary Data Set S8). This suggests that FaRIF, in addition to regulating the expression of ABA-related genes, may stabilize their mRNAs in conjunction with FaYTH proteins, integrating transcriptional and post-transcriptional regulation during ripening.

Proteins involved in protein folding and processing were also identified as FaRIF interactors. Among them, the chaperonin complex TriC/CCT subunit CCT3 was validated as an interactor (Fig. 7C). The TriC/CCT complex assists in folding specific eukaryotic proteins (Spiess et al., 2004), and has been reported to directly regulate HSF1, a stress-responsive TF (Neef et al., 2014). Whether the interaction between FaRIF and CCT3 serves as ripening-specific function remains to be elucidated.

Finally, two Ca^2+^-dependent lipid-binding proteins containing a single C2 domain, FaCalB1 and FaCalB2, were identified, with FaCalB1 validated by CoIP (Fig. 7C). Calcium acts as an intracellular messenger in ABA signaling (Schroeder et al., 2001) and stimulates flavonoid-related genes and anthocyanin accumulation in strawberry (Xu et al., 2014). Single C2 domain proteins like CalB1 Arabidopsis ortholog, and a close related protein in rice, *OsERG1*, are involved in plant defense signaling (Cooper et al., 2003; Kim et al., 2003). Interestingly, differences in the membrane binding capability and subcellular localization have been found in close homologs of FaCalB1 in rice (Kim et al., 2003; Kang et al., 2011) and barley (Ouelhadj et al., 2006), being the latter localized in nuclei as FaCalB1 (Fig. 7B). It is intriguing to speculate that FaCalB proteins mediate calcium-mediated ABA signaling and the ripening progress in coordination with FaRIF, which requires further investigation.

Future studies should define the co-regulatory mechanisms and functional implications of FaRIF’s interactions, particularly in coordination with other ripening-related NAC TFs, as well as in mRNA stability and post-transcriptional regulation, and in protein homeostasis.

## Material and Methods

### Plant material and growth conditions

Stable *35Spro:FaRIF-RNAi*, *35Spro:FaRIF* and *35Spro:FaRIF-GFP* transgenic lines were previously established in the group as described in (Martín-Pizarro et al., 2021). The full-length *FaRIF* constructs correspond to the *FaRIF(3B)* ortholog. *F.* × *ananassa* cv. Camarosa control and transgenic strawberry adult plants were grown and maintained in a shading house (IFAPA, Churriana, Málaga, Spain) and greenhouse (IHSM, Málaga, Spain) conditions.

### Transcriptome analysis by RNA-seq

The RNA-seq data previously generated from RNAi lines of *FaRIF* and mapped using the assembly and annotation version v4.0.a1 of the *F. vesca* reference genome (Martín-Pizarro et al., 2021) were remapped against the *F.*× *ananassa* cv. Camarosa genome assembly v1.0.a1 (Edger et al., 2019), and annotated using *F.* × ananassa cv. Camarosa reference genome v1.0.a2 (Liu et al., 2021). This analysis was conducted using the European Galaxy server (https://usegalaxy.eu/) (Goecks et al., 2010). Sequence alignment was performed using the HISAT2 tool for sequence alignment, followed by read counting with the htseq-count tool. The edgeR tool was employed for differential expression analysis of count data. MapMan bins were used for the assignment of functional categories to the differentially expressed genes with more than two-fold down- or upregulation in the RNA lines at each ripening stage (Log_2_(FC) ≤ -1 or ≥ 1; FDR ≤ 0.05) (Schwacke et al., 2019).

### Protein structure prediction

Protein sequences of FvRIF and FaRIF were used to predict their structure using AlphaFold 3 (Abramson et al., 2024). The five models predicted were visualized and edited using PyMOL software (Schrödinger and DeLano 2020).

### Chromatin Immunoprecipitation assay

The *35Spro:RIF:GFP* stable transgenic lines and wild-type plants (Martin-Pizarro et al., 2021) were used for ChIP, following a modified protocol from Gendrel et al. (Gendrel et al., 2005). Fixation was carried out using 1 g of ground powder without vacuum using extraction buffer 1 (0.4 M sucrose, 10 mM HEPES pH 7.5, 0.1 mM phenylmethanesulfonyl fluoride (PMSF), 10 mM MgCl_2_, 1% PVP-40, 5 mM β-ME, complete protease inhibitor cocktail and 1% formaldehyde) at room temperature inverting the tubes. To stop cross-linking 0.15 M final concentration of fresh glycine was added, and the tubes were incubated and inverted at room temperature for 10 min. The samples were filtered through Miracloth (Millipore) and centrifuged. The pellets were resuspended in extraction buffer 2 (0.4 M sucrose, 10 mM HEPES pH 7.5, 0.1 mM PMSF, 10 mM MgCl_2_, 1% PVP-40, 1% Triton X-100, 5 mM β-ME, complete protease inhibitor cocktail). Then, extraction buffer 3 (1.7 M sucrose, 10 mM HEPES pH 7.5, 0.1 mM PMSF, 20 mM MgCl_2_, 1% PVP-40, 0,15% Triton X-100, 5 mM β-ME, complete protease inhibitor cocktail) was used. The isolated nuclei were transferred to nuclei buffer lysis (50 mM Tris-HCl, 10 mM EDTA, 1% SDS, complete protease inhibitor cocktail) and sonicated using BioruptorTM UCD-200 sonicator (Diagenode). The chromatin was sheared into fragments from 200 bp to 1000 bp. The supernatant was transferred to a new tube for immunoprecipitation, and a small aliquot was kept as the “input DNA control”. The chromatin was diluted 10 times with ChIP dilution buffer (20 mM Tris-HCl, 2 mM EDTA,150 mM NaCl, 0.1% SDS, 1% Triton X-100) and pre-cleared with 50 µL protein A–agarose beads (Santa Cruz Biotechnology) for 1 h on a rotator at 4 °C. The samples were centrifuged to remove the beads, and the chromatin was incubated with anti-GFP antibody (Abcam; ab290) overnight. The chromatin was then incubated with protein A–agarose beads for 2 h on a rotator at 4 °C. The beads were washed low-salt wash buffer (20 mM Tris-HCl pH 8, 2 mM EDTA, 150 mM NaCl, 0.1% SDS, 1% Triton X-100, high-salt wash buffer (20 mM Tris-HCl pH 8, 2 mM EDTA, 500 mM NaCl, 0.1% SDS, 1% Triton X-100), LiCl wash buffer (10 mM Tris-HCl pH 8, 1 mM EDTA, 250 mM LiCl, 1% sodium deoxycholate, 1% NP-40 substitute), and finally TE buffer (10 mM Tris-HCl pH 8, 1 mM EDTA).To elute the complex DNA-beads, the samples were treated with elution buffer (1% SDS, 0.168 g NaHCO_3_), vortexed and incubated at 65 °C for 15 min in agitation at 700 rpm. The reverse cross-linking was conducted by adding 5 M NaCl and incubating overnight at 65 °C the immunoprecipitated and the input samples. The samples were treated with 1 μL of RNase A/T1 (10 mg/mL) at 37 °C. Subsequently, 20 μL 1M Tris-HCl pH 6.5, 10 μL 0.5 M EDTA, and 2 μL Proteinase K were added and incubated at 55 °C for 1 h. Another 1 μL of Proteinase K was added and incubated at 37 °C for 1 h. The immunoprecipitated and input DNA were purified with MinElute Reaction Cleanup kit (QIAGEN). The resulting samples were analyzed by qPCR to confirm enrichment using primers for primers of *FaRIF, FaNAC042*, and *FaPL2* (Supplementary Data Set S9). Two independent biological replicates from the immunoprecipitated and input DNA control were prepared for libraries and high throughput sequencing by Illumina platform Novaseq6000 according to the manufacturer’s instructions.

### ChIP-seq analyses

DNA libraries were sequenced in an Illumina NovaSeq600 system (Novogene, Tianjin, China), and raw reads (150 nucleotides, paired ends) were trimmed of adapter sequences and quality filtered with Trim Galore! (https://github.com/FelixKrueger/TrimGalore), using default parameters (–q 20 – e 0.1 –length 20). Clean reads were mapped against the *F.*× *ananassa* cv. Camarosa reference genome assembly v1.0.a1 (Edger et al., 2019) and annotation v1.0.a2 (Liu et al., 2021) with Bowtie2 and default parameters (Langmead and Salzberg, 2012). Peak calling was performed with MACS2 (Feng et al., 2012) using an effective genome size of 740 Mb and treating both replicates independently. Significant peaks were considered when enriched over two-fold in the immunoprecipitated sample relative to their corresponding input sample and FDR-adjusted p-value lower than 0.001. Trimming, mapping, and peak calling steps were performed in the European Galaxy server (https://usegalaxy.eu/) (Goecks et al., 2010).

Significant peaks overlapping in the two replicates were obtained with BEDTOOLS intersect (Quinlan and Hall, 2010) and these were considered as FaRIF-bound fragments for further analysis. Peaks were annotated relative to the nearest gene and the TSS with the R package ChIPseeker (Yu et al., 2015), using the annotation file Fragaria_ananassa_v1.0.a2.genes.gff3. Enriched binding sites were identified using the MEME-ChIP tool in MEME Suite (Bailey et al., 2009), using as input 1,000 bp centered at the summits of the 13,620 significant peaks (fold-enrichment > 2; q-value < 0.001).

### Analysis of peak distribution enrichment in the different *F.* × *ananassa* subgenomes

To determine the binding preferences of FaRIF for the different subgenomes of *F.* × *ananassa*, a Chi-Square analysis for each of the seven chromosomes, including the four homoeologous chromosomes from the different subgenomes, was conducted. The observed number of peaks was based on either the total number of peaks or those located within the region spanning 2 kb upstream to 100 bp downstream of the TSS, which were consistently found in both biological replicates. The expected numbers were calculated based on the total size of the specific chromosome, including all four subgenomes. A difference was considered statistically significant when the Chi-Square p-value was ≤ 0,01. A 75th percentile threshold was applied to further identify which subgenomes contributed to the difference.

### TurboID plasmid construction

The coding sequence of *FaNAC035* (*FaRIF*) was cloned into two different vectors using the Ti-TAN plasmid toolbox (Tan et al., 2024). To confirm proper nuclear localization, the *FaRIF* CDS was fused to the *TurboID* (*TbID*) and *GFP* CDSs at the C-terminal region, generating pCM44 (*35Spro:RIF-TbID-GFP)*. For the final TbID experiment, the *FaRIF* CDS was fused to the 3xHA-TbID fusion protein at the C-terminal, resulting in pCM45 (*35Spro:FaRIF-3xHA-TbID*). A *35Spro:3xHA-TbID-3xNLS* vector, containing TurboID with three tandem copies of nuclear localization, was included as a negative control. The primers for the plasmids construction are listed in Supplementary Table S9.

### Transient expression in strawberry fruits

Fruits were labeled and agroinfiltrated at the white stage of fruit development with *Agrobacterium tumefaciens* strain GV3101 carrying pCM45, or the negative control vector and p19, as previously described (Hoffmann et al., 2006). The fruits were agroinfiltrated with the suspension, and when they began to ripen, they were infiltrated with a solution containing 50 µM of biotin and 2.5 mM MgCl_2_ or only with 2.5 mM MgCl_2_ (mock). After 5 hours, the fruits were harvested, immediately frozen in liquid nitrogen, deachened, and ground into a fine powder. Western blot analysis was used to confirm recombinant protein expression using an α-HA antibody, and the enrichment of biotinylated protein was validated with streptavidin-HRP. The experiment was performed by obtaining three independent pools for construction, each composed of seven independent fruits (n∼21 per construct).

### Nuclear protein extraction and western blot analysis

Nuclear protein extraction was performed as previously described by Bouyer et al (2011). Proteins were separated on 10% SDS-PAGE and transferred onto an Immobilon-P PVDF membrane (Millipore, Catalog number IPVH00010) using the Trans-Blot Turbo Transfer System (Bio-Rad). The membranes were blocked using 5% fat-free milk in TTBS for α-HA antibody (Sigma-Aldrich, Catalog number H3663) and 3% Bovine serum albumin (BSA) in TTBS for streptavidin-HRP (Abcam, Catalog number ab7403). After blocking for 1h at room temperature, membranes were washed with TTBS and incubated with the respective antibodies for 2 hours or overnight. After incubation, membrane with α-HA antibody was washed three times with TTBS and incubated with the secondary antibody anti-mouse IgG-Peroxidase (Sigma-Aldrich, Catalog number A4416). Signal detection was performed using Clarity ECL Western Blotting Substrates (Bio-Rad). Equal protein loading was confirmed by staining membranes with Coomassie Brilliant Blue (CBB) R 250.

### Samples preparation for mass spectrometry

To remove the free biotin in the nuclear protein samples, the desalting columns (PD-10 Cytiva 17-0851-01) were used. First, the columns were equilibrated using 5 mL of the equilibrated solution: 100 mM Tris-HCl (pH 7.5), 150 mM NaCl, 10% Glycerol, 5 mM EDTA, 2 mM DTT, three times. Subsequently, 2 mL of the nuclear protein extraction was passed through the column. The tubes were centrifuged at 1000g, 4 °C for 2 min, and the protein elution was collected and transferred to a new 2 mL tube. To enrich biotinylated proteins from the samples, 80 µL of streptavidin-coated magnetic beads (Dynabeads™ MyOne™ Streptavidin C1, Invitrogen, catalog number 65001) per sample were washed twice with extraction buffer (100 mM Tris-HCl pH 7.5, 150 mM NaCl, 10% Glycerol, 5 mM EDTA, 5 mM DTT, 1% protease inhibitors cocktail, 1 mM PMSF, 1% NP40) at room temperature. The samples were incubated with the pre-cleaned beads for 4 hours on a rotor at 4 °C. After the incubation, the magnetic beads were washed with different buffers: 1.7 mL Buffer I (0.5% SDS in water), 1.7 mL Buffer II (50 mM HEPES pH 7.5, 500 mM NaCl, 1 mM EDTA, 1% Triton-X-100, 0.5% Sodium-Deoxycholate), 1.7 mL Buffer III (10 mM Tris pH 7.5, 250 mM LiCl, 1 mM EDTA, 0.5% NP-40, 0.5 % Sodium-Deoxycholate). The beads were washed twice with 1.7 mL of 50 mM Tris-HCl pH 7.5 to remove the detergent. This was followed by three washes with 1X PBS buffer at room temperature. To confirm the successful enrichment of the samples, 80 µL was taken for Western Blot analysis. The rest of the PBS buffer was removed and the beads were flash-frozen and kept at -80°C until they were sent to the LC-MS/MS analysis.

### NanoLC-MS/MS analysis and MS data processing

The biotinylated proteins were eluted from streptavidin beads and purified with a 12% NuPAGE Novex Bis-Tris Gel (Invitrogen). Tryptic in-gel digestion of proteins was performed as described previously (Borchert et al., 2010) and the extracted peptides were desalted using C18 StageTips (Rappsilber et al., 2007). The eluted peptides were subjected to LC-MS/MS analysis. Peptide analysis was conducted on a Proxeon EASY-nLC 1200 system coupled to an Orbitrap QExactiveHF mass spectrometer (Thermo Fisher Scientific), peptides were eluted with a 49 min segmented gradient of 10–33-50-90% HPLC solvent B, 80% acetonitrile (ACN) in 0.1% formic acid (FA).

The MS raw data were processed with MaxQuant software suite version 2.2.0.0 (Cox & Mann, 2008). Database search was performed using the Andromeda search engine (Cox et al, 2011), a module of the MaxQuant. The processing was done by searching MS/MS spectra with *Fragaria* × *ananassa* database with 1,004 entries. The FaRIF-TurboID sequence and a database consisting of 286 commonly observed contaminants were added to the search. iBAQ values (Schwanhäusser et al., 2011) were LFQ settings selected and matched between runs enabled only within triplicates. In the database search, full tryptic specificity was required and up to two missed cleavages were allowed. Carbamidomethylation of cysteine was set as fixed modification, whereas oxidation of methionine and acetylation of protein N-terminus were set as variable modifications. Mass tolerance for precursor ions was set to 4.5 ppm and for fragment ions to 20 ppm. Peptide, protein and modification site identifications were reported at an FDR of 0.01, estimated by the target/decoy approach (Elias and Gygi, 2007). For protein group quantitation, a minimum of two quantified peptides were required. Statistical analysis was performed with the Perseus software suite (version 2.0.5.0.). Reverse hits and potential contaminants were filtered out from the protein dataset, along with those identified only by site. Median and mean values for each group were calculated with respective standard deviations. A new categorical column was created to specify the unique entries for each group. Data was normalized by log2 transformation, Student’s *t*-test comparison was used to identify significantly changing entries between “C1” and “Positive” samples with a permutation-based FDR (s0 = 0 and FDR ≤0.05), for the t-test data was filtered for at least three valid values in at least one group.

### Subcellular localization

To validate the *FaRIF* interacting proteins obtained from the TurboID, five candidate proteins were selected for transient expression and subcellular colocalization using *Nicotiana benthamiana*: *FaNAC035* (*FaRIF*), *FaNAC021*, *FaNAC034*, *FaCalB1* and *FaCCT-3*. The coding sequence was amplified from cDNA prepared from red fruits of the octoploid *F.* × *ananassa* cv. Camarosa with gene-specific primers. All of them were cloned into pGWB554, a binary vector fused to the red fluorescent protein (mRFP) at the C-terminal region and driven by the *35S* constitutive promoter. To generate *35Spro:FaRIF-GFP*, the full CDS was amplified from the same cDNA and cloned into the pK7WG2D binary vector generating pCM24. The empty vector, pGWB554, was used as a control (*35Spro:mRFP*). The vectors were transferred to *Agrobacterium tumefaciens* (strain GV3101). The bacteria were then mixed and infiltrated at an OD_600_ of 0.4 for the constructs with mRFP, 0.4 for pCM24 and 0.2 for p19 in 3 to 4 weeks-old *N. benthamiana* leaves. The plants were maintained in chambers in long-day conditions (16 h light/8 h dark) controlled photoperiod at 23 °C. GFP and mRFP fluorescence signals were observed 2 days post-infiltration and the images were taken with a ZEISS LSM 880 confocal laser-scanning microscope with 40x objective. Scan setting, excitation at 488 nm and emission 500-550 nm for GFP and excitation at 543 nm and emission 600-650 nm for RFP. The primers for the plasmids construction are listed in Supplementary Table S9.

### Coimmunoprecipitation assay

The CoIP experiments were conducted using the coinfiltrated *N. benthamiana* leaves. Approximately 1 g of leaves were ground to a fine powder in liquid nitrogen. The total protein extraction was conducted with 2 mL extraction buffer (100 mM Tris-HCl pH 7.5, 150 mM NaCl, 10% glycerol, 5 mM EDTA, 10 mM DTT, 1% protease inhibitor cocktail (P9599, Sigma-Aldrich), 1% NP-40, 10 mM Na_2_MoO_4_, 10 mM NaF, 2 mM Na_3_VO_4_ and incubated on a rotor at 4 °C for 30 min. Samples were centrifuged at 14,000 rpm at 4 °C for 20 min. Then, the supernatant was incubated with GFP-Trap magnetic agarose beads (Chromotek) at 4 °C for 2 h. The beads were collected and washed four times with wash buffer (100 mM Tris-HCl pH 7.5, 150 mM NaCl, 10% glycerol, 2 mM DTT, 1 % protease inhibitor cocktail (P9599, Sigma-Aldrich), 0.2 % NP-40, 10 mM Na_2_MoO_4_, 10 mM NaF, 2 mM Na_3_VO_4_). Proteins bound to the beads were eluted in 100 µL of Laemmli buffer (2X) at 95 °C for 10 min. The experiment was checked by immunoblotting using anti-RFP or anti-GFP antibodies.

### Dual-Luciferase reporter assay

One *N. benthamiana* leaf from six independent 3- to 4-week-old plants were infiltrated on the abaxial side using a 1 mL syringe containing an agroinfiltration solution (10 mM MES, pH 5.6, 10 mM MgCl2, and 1 mM acetosyringone), each used as an independent biological replicate. Promoter sequences were cloned into the pGreenII 0800-LUC plasmid (Hellens et al.,2005) One or two TF constructs were co-infiltrated at a final OD_600_ of 0.6 and their corresponding putative target promoter at a final OD_600_ of 0.4. In all mixtures, an *A. tumefaciens* clone overexpressing the RNA silencing suppressor p19 was co-infiltrated at an OD_600_ of 0.2. pK7WG2 empty vector was used as the negative control. Two 1 cm² leaf fragments were analyzed per inoculation. Firefly luciferase and Renilla luciferase activities were measured 3 days after infiltration using the Dual-Luciferase Reporter Assay System (Promega) and the Glomax® Navigator System (Promega) luminometer. Data are expressed as the firefly luciferase/renilla (LUC/REN) ratio. The primers for the plasmids construction are listed in Supplementary Table S9.

## Supporting information

Supplementary Data Set S1

Supplementary Data Set S2

Supplementary Data Set S3

Supplementary Data Set S4

Supplementary Data Set S5

Supplementary Data Set S6

Supplementary Data Set S7

Supplementary Data Set S8

Supplementary Data Set S9

## Author Contributions

C.M.-P., G.Q., and D.P. conceived and designed the research. C.M.-P. performed and analyzed most experiments and data. M.F.P. performed the transactivation assays in Nicotiana. J.M.F.-Z. analyzed the ChIP-seq data. R.L.-D. contributed to the TurboID-based proximity labeling experiment. C.M.-P. and D.P. secured funds, designed and supervised the experiments, and wrote the article. All authors commented on the article.

## Acknowledgments

This work was supported by a grant from the Spanish Ministry of Science and Innovation and Universities (MICIU, PID2021-123677OB-I00 to D.P., and PID2023-149400NB-I00 to J.M.F.-Z.) and funding from the Junta de Andalucía (UMA20-FEDERJA-093 and Postdoctoral program, POSTDOC_21_00893 to C.M.-P.), EMBO (EMBO Scientific Exchange Grant, ref 10186 to C.M.-P.), MICIU (Juan de la Cierva Program, JDC2022-048407-I to M.F.-P.), and by the Excellence Strategy of the German Federal and State Governments (to R.L.-D.). We thank Dr. José F. Sánchez Sevilla for facilitating growing the transgenic plants at the Andalusian Institute for Research and Training in Agriculture, Fishery, Food and Ecological Production (IFAPA) facilities in Churriana, Málaga, Spain, and Francisco Durán for its maintenance, respectively. We thank GeminiTeam (ZMBP, University of Tübingen) for their support with the TurboID experiments, and Irina Droste Borel (Proteome Center Tübingen (PCR), University of Tübingen), for the NanoLC-MS/MS analysis and MS data processing.

## Figures

**Supplementary Fig. S1.**
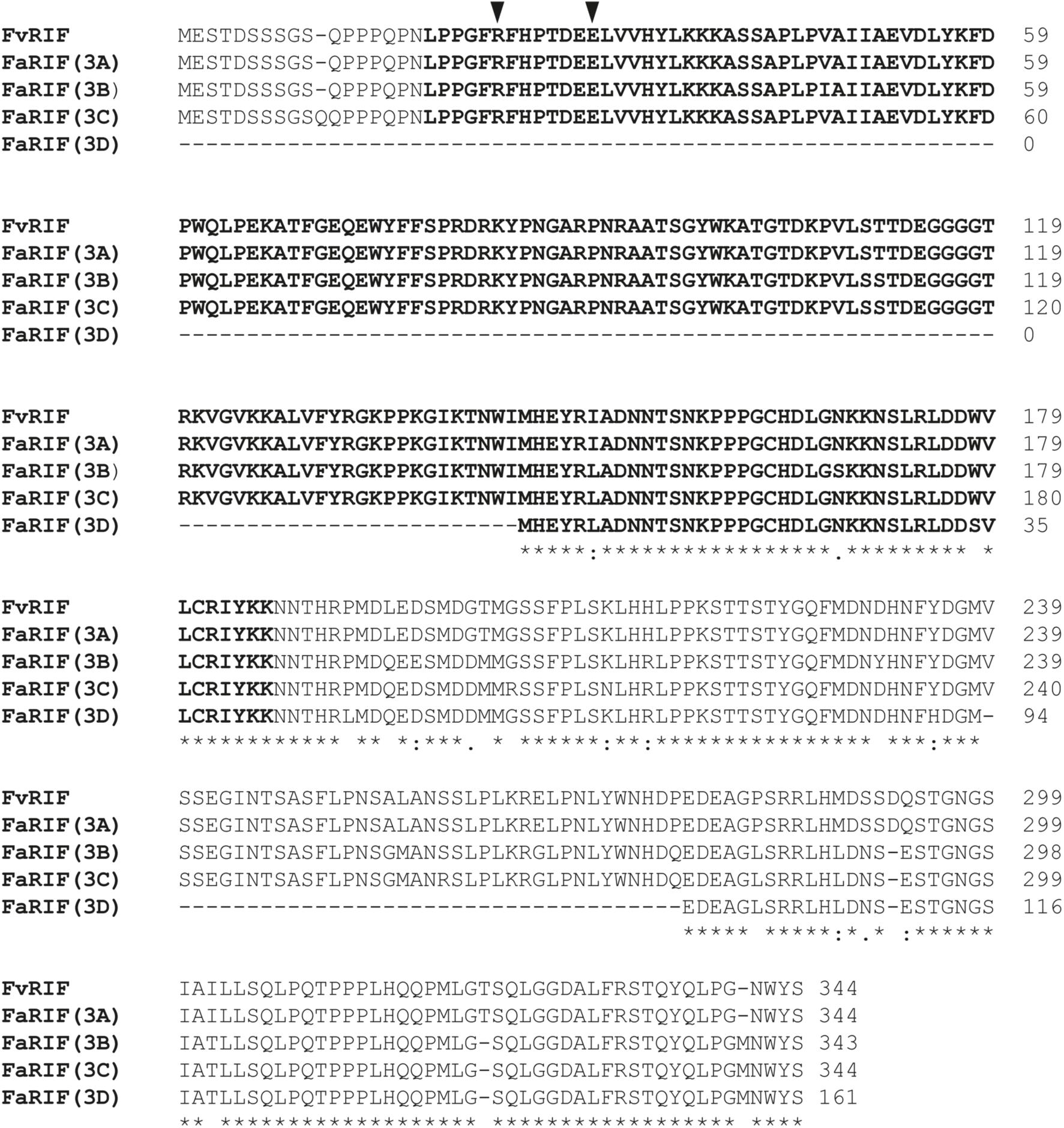
Alignment of RIF proteins from *F. vesca* and *F.* × *ananassa*. The NAC domain is depicted in bold fonts. Triangles denote conserved residues involved in the dimerization of NAC proteins, as predicted in our analysis using Alphafold 3 (Abramson et al., 2024) and by Ernst and collaborators (Ernst et al., 2004).

**Supplementary Fig. S2.**
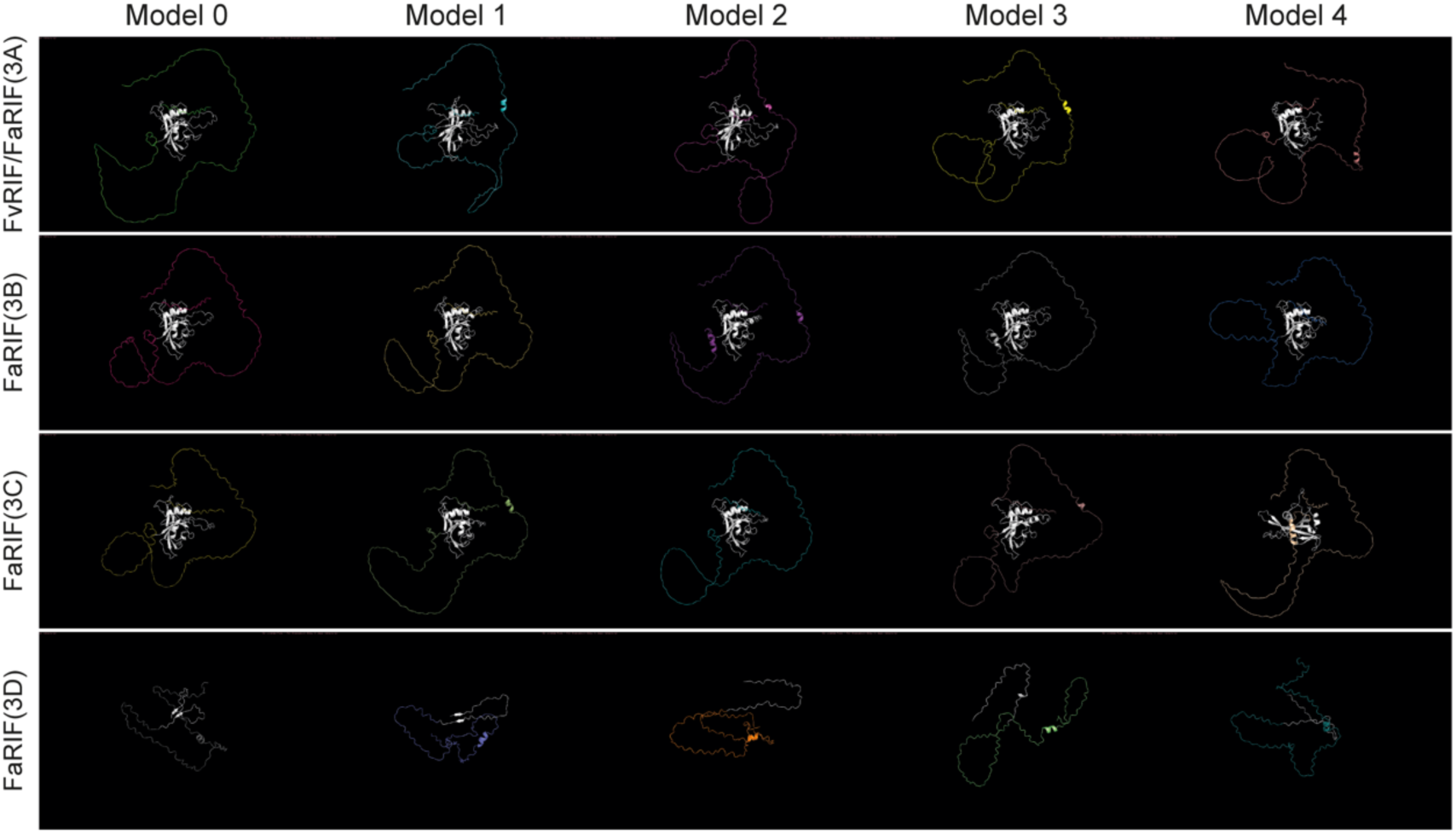
Protein structure prediction of FvRIF and FaRIFs proteins using Alphafold 3 (Abramson et al., 2024). The five models predicted were aligned against model 0 of FvRIF/FaRIF(3A) protein using PyMol. The NAC domain is represented in white color.

**Supplementary Fig. S3.**
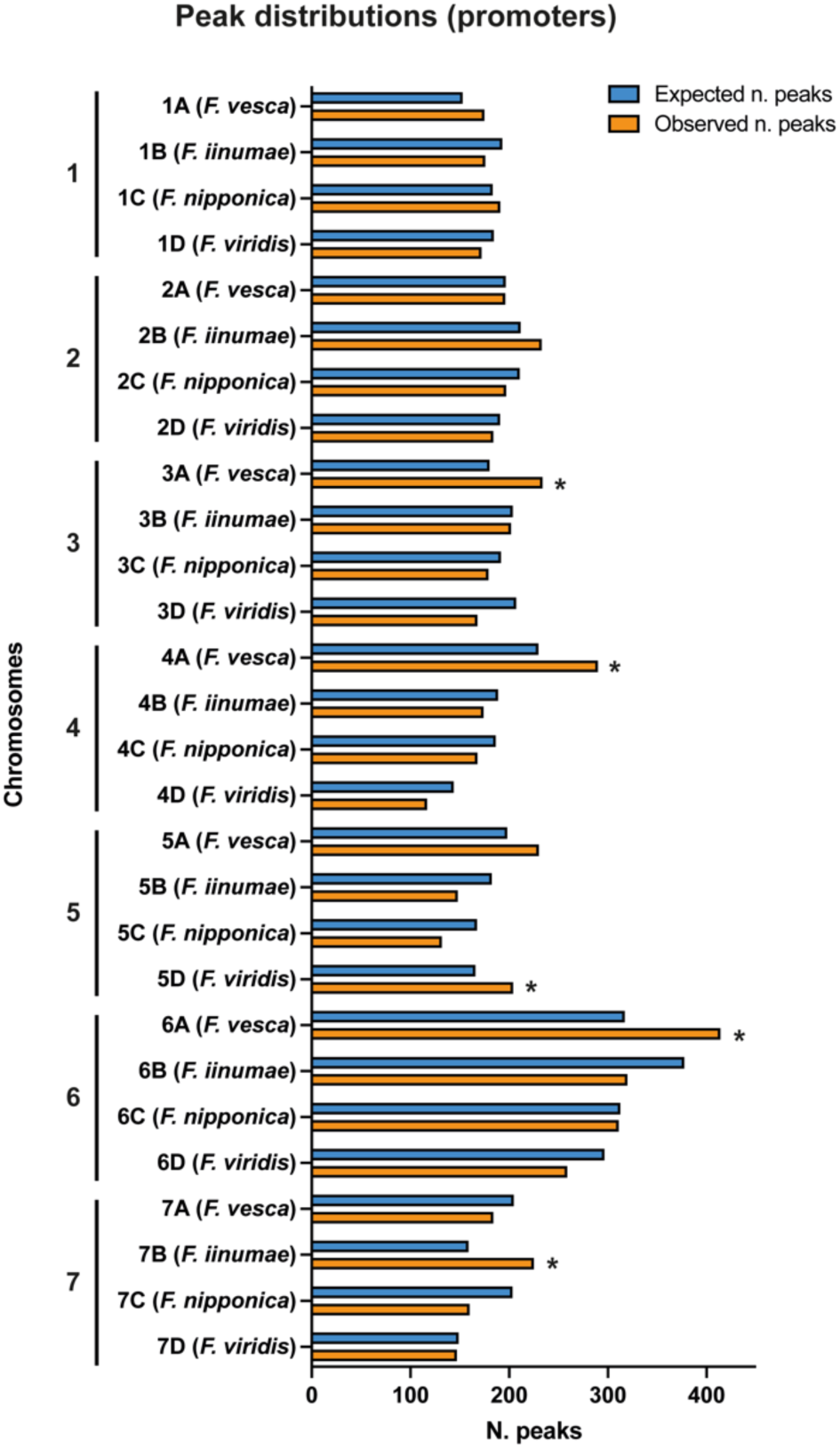
Genome-wide distribution of FaRIF binding sites across all *F.* × *ananassa* chromosomes. Number of observed (orange bars) and expected (blue bars) binding sites for those peaks located within the region spanning 2 kb upstream to 100 bp downstream of the TSSs, also consistently identified in both biological replicates. The expected number of binding sites was calculated by dividing the total number of peaks for each chromosome set by the total size of that set. Asterisks (*) indicate statistically significant differences between observed and expected binding sites.

**Supplementary Fig. S4.**
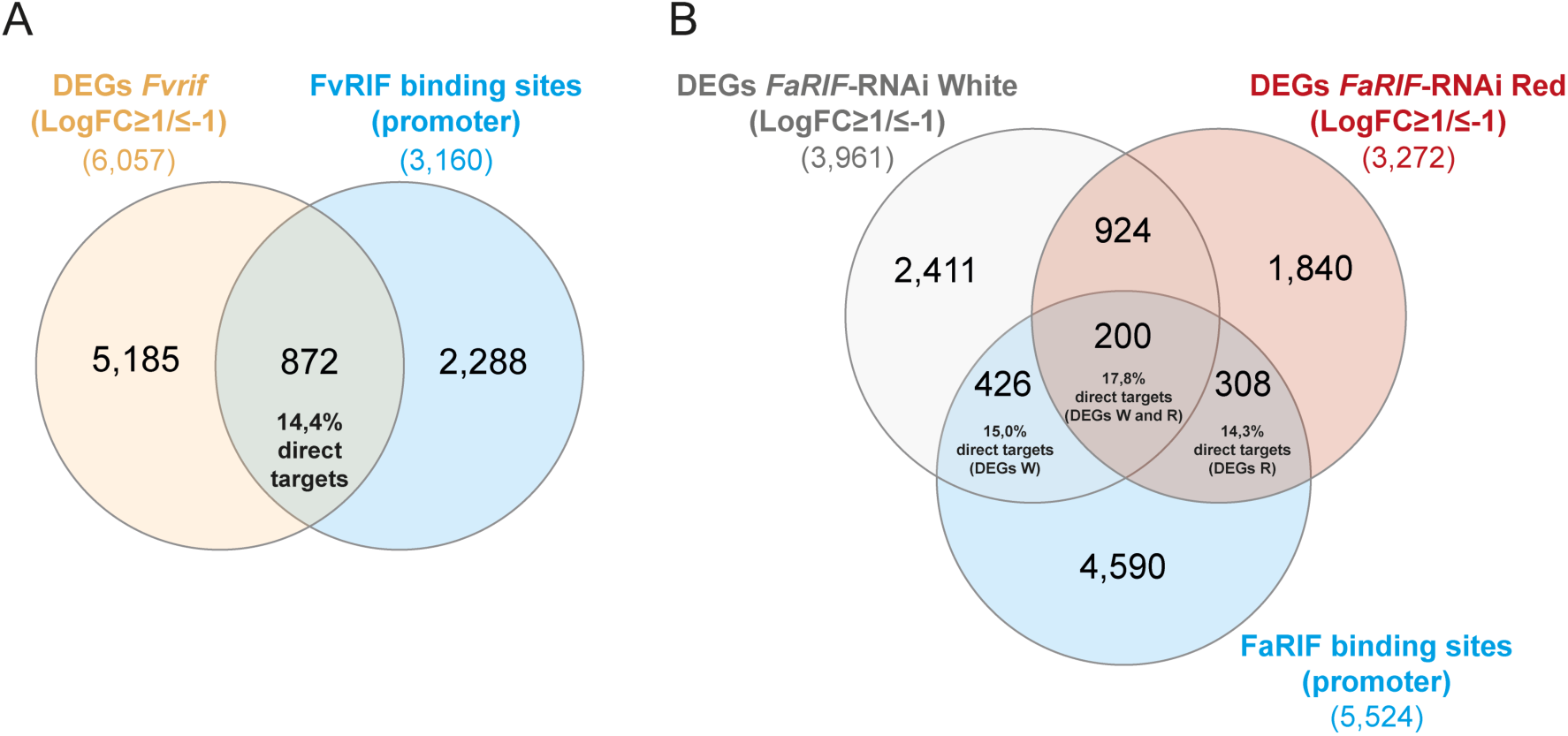
Comparison of RNA-seq and ChIP/DAP-seq data to identify potential direct targets of RIF in *F.* × *ananassa* and *F. vesca.* **A)** Comparison of all the DEGs with at least two-fold change (up- or downregulation) identified in *Fvrif* mutant lines with FvRIF binding sites located in the promoter regions, using data from Li and collaborators (Li et al., 2023). **B)** Stage-specific comparison of DEGs with at least two-fold change (up- or downregulation) identified in *FaRIF*-RNAi lines at each ripening stage with FaRIF binding sites in the promoter regions.

**Supplementary Fig. S5.**
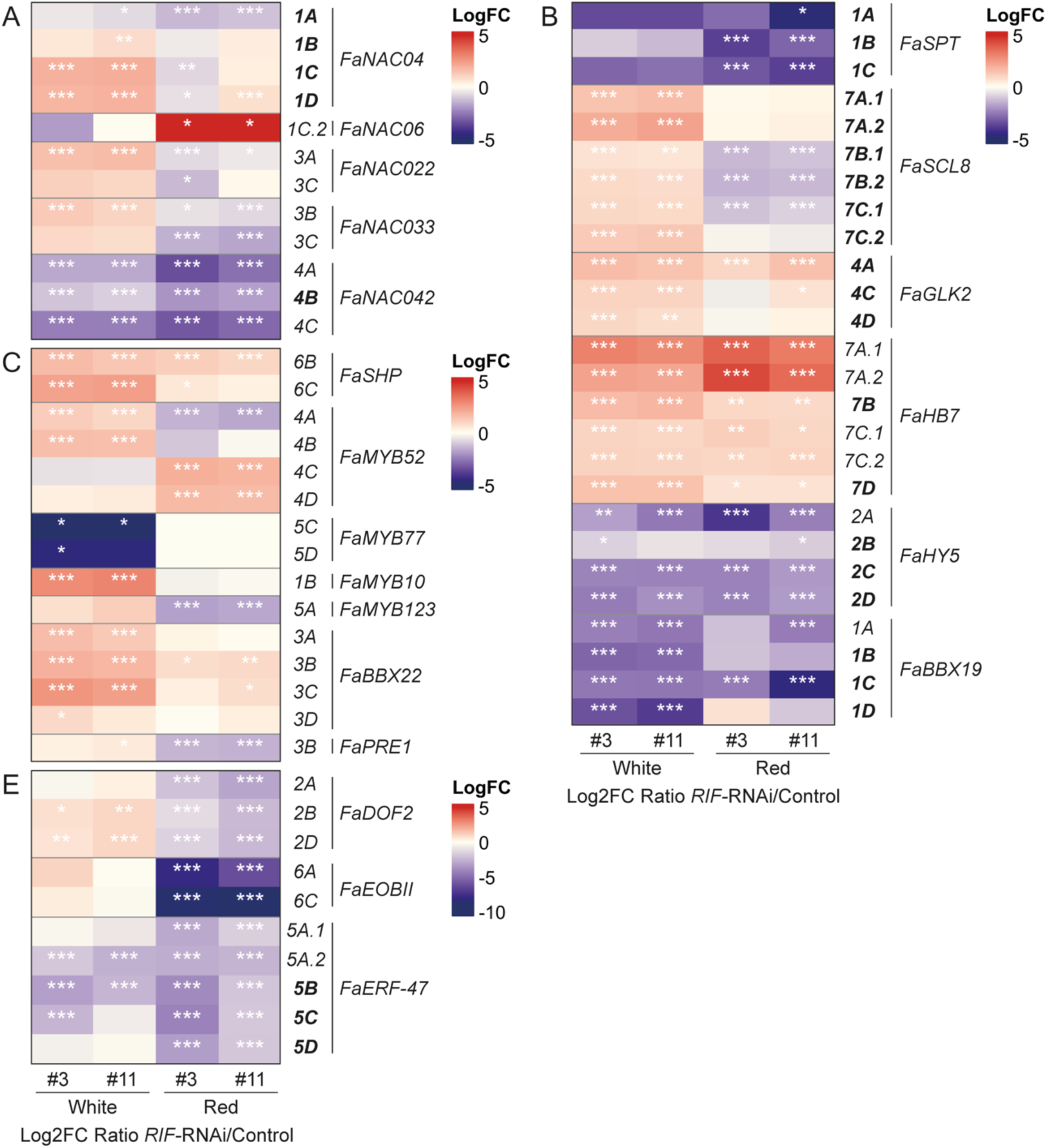
Transcription factors directly and indirectly regulated by FaRIF. **B) to E)** Heatmaps displaying the expression of FaRIF-regulated transcription factors, with direct targets in bold. Panels include transcription factors from the NAC family **(A)**, involved in fruit development and ripening-related processes **(B)**, and in anthocyanin **(C)** and aroma compounds **(D)** biosynthesis. Significant DEGs are marked with asterisks (***FDR≤0,0005; **FDR≤0,005; *FDR≤0,05). Detailed information is available in Supplementary Data Set S7.

**Supplementary Fig. S6.**
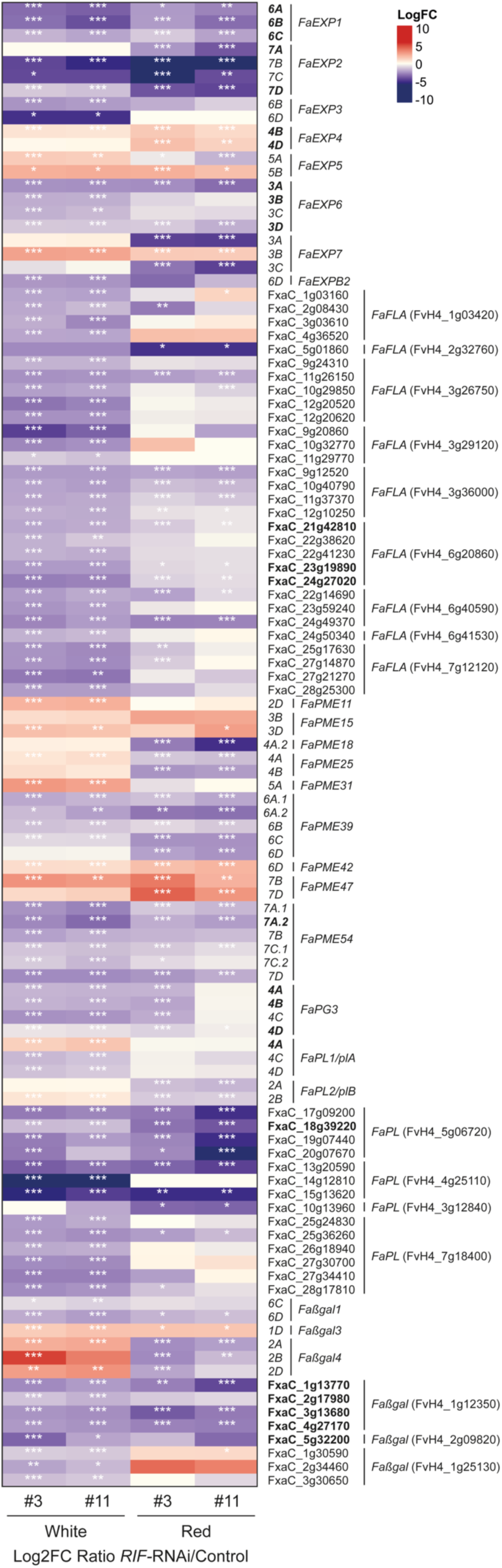
Cell wall-related genes directly and indirectly regulated by FaRIF. Heatmap displaying the expression of FaRIF-regulated cell wall modifying genes, with direct targets in bold. Significant DEGs are marked with asterisks (***FDR≤0,0005; **FDR≤0,005; *FDR≤0,05). Detailed information is available in Supplementary Data Set S7.

**Supplementary Fig. S7.**
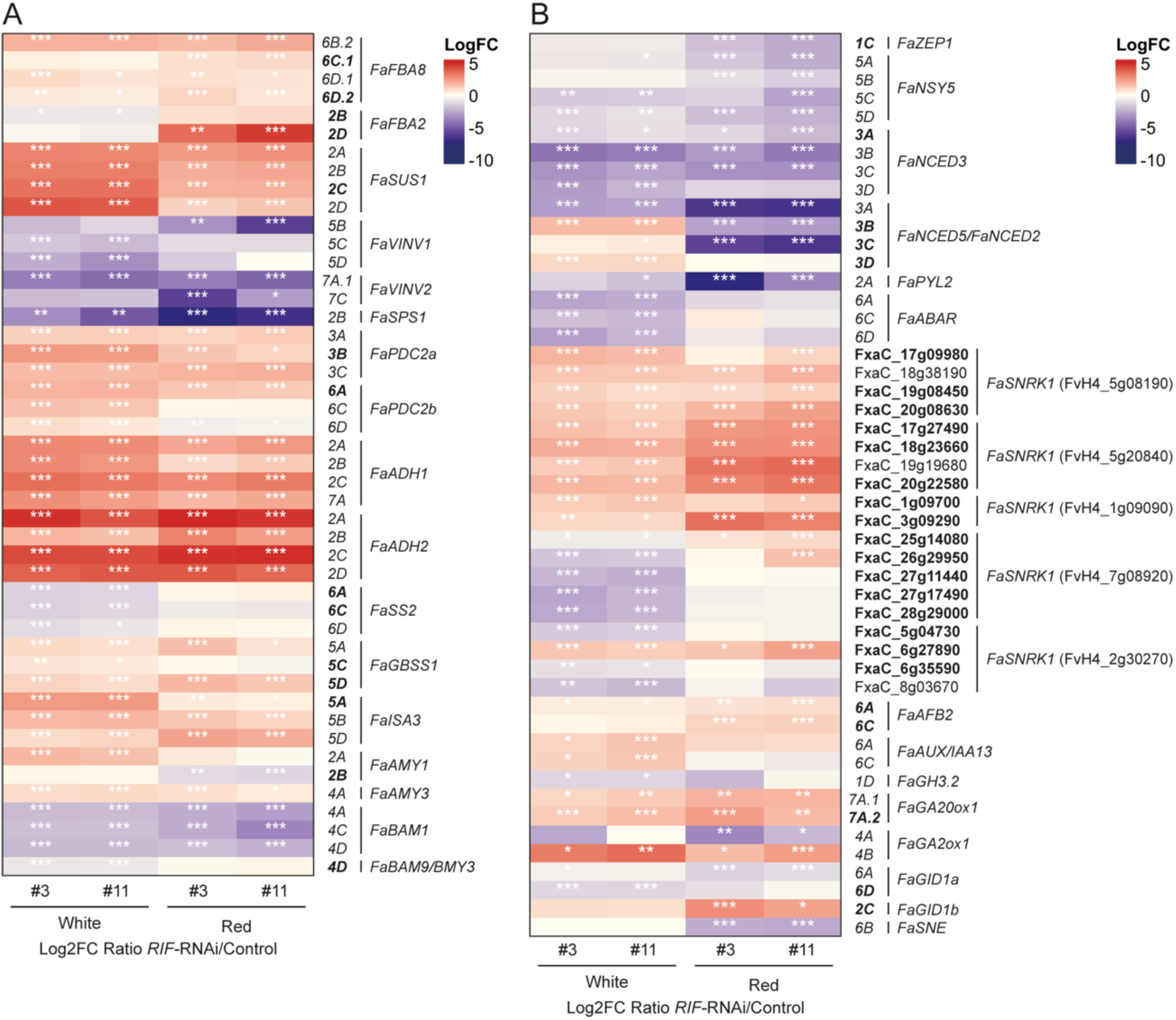
Carbohydrate metabolism and hormone-related genes directly and indirectly regulated by FaRIF. Heatmaps displaying the expression of FaRIF-regulated genes related to carbohydrate metabolism **(A)**, and hormone biosynthesis and signaling pathways **(B)**, with direct targets in bold. Significant DEGs are marked with asterisks (***FDR≤0,0005; **FDR≤0,005; *FDR≤0,05). Detailed information is available in Supplementary Data Set S7.

**Supplementary Fig. S8.**
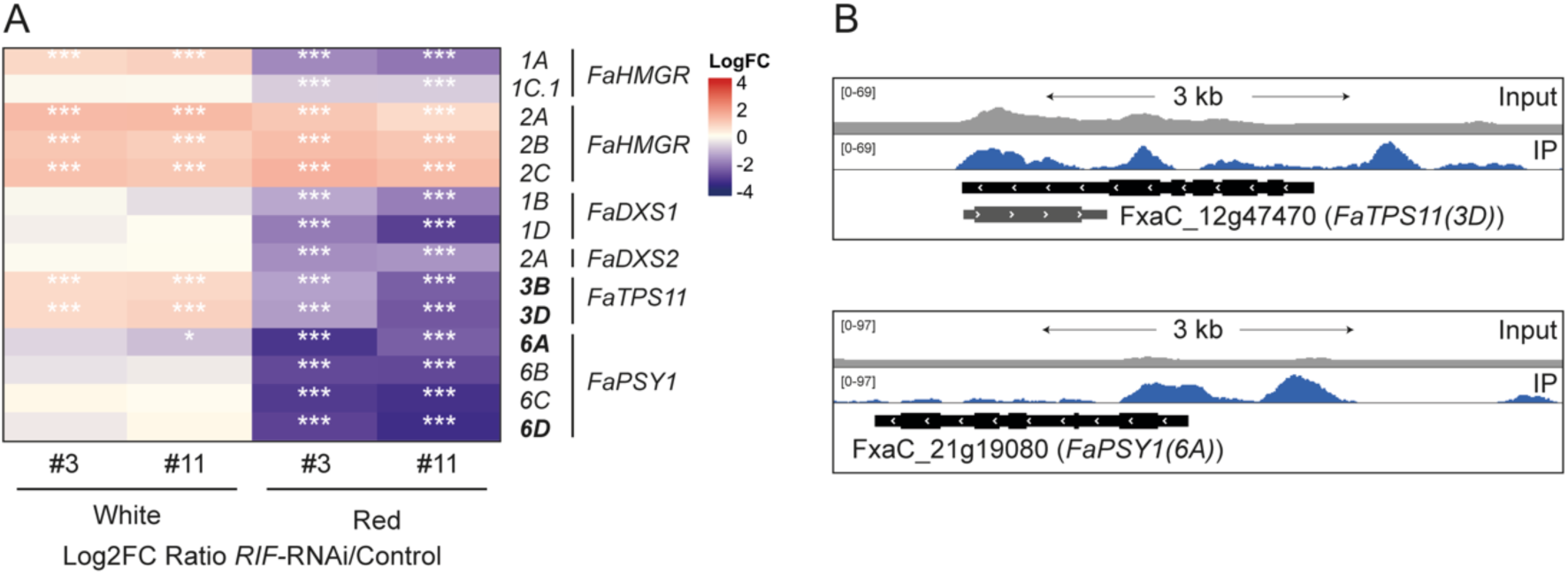
Terpenoid biosynthetic genes directly and indirectly regulated by FaRIF. **A)** Heatmap displaying the expression of FaRIF-regulated genes related to terpenoid metabolism. Significant DEGs are marked with asterisks (***FDR≤0,0005; **FDR≤0,005; *FDR≤0,05). **B)** ChIP-seq peaks showing FaRIF binding sites and input reads over representative homoeologs of direct targets involved in terpenoid biosynthesis. Top panel includes gene FxaC_12g47480 depicted in gray. Detailed information is available in Supplementary Data Set S7.

**Supplementary Fig. S9.**
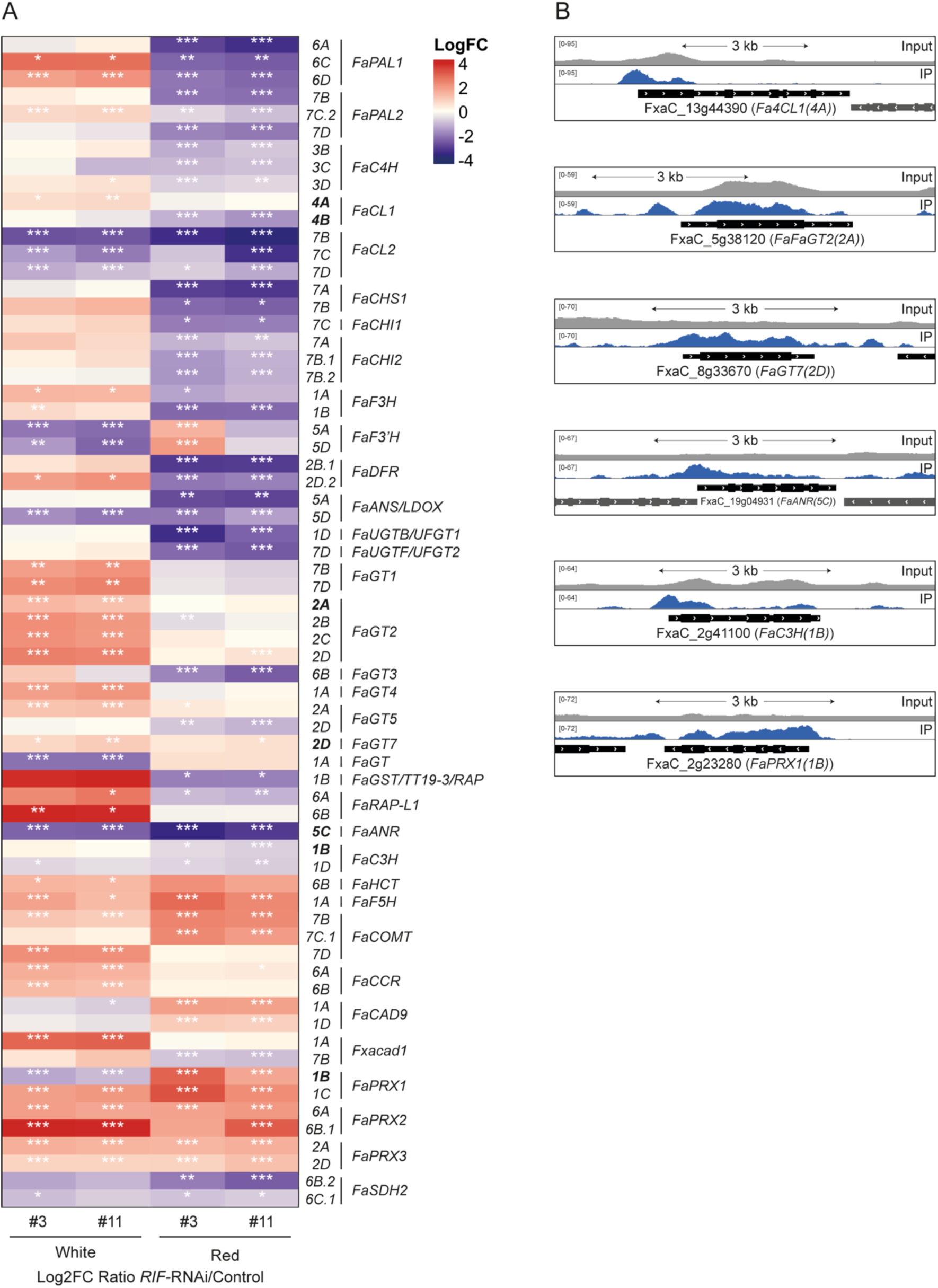
Phenylpropanoid pathway biosynthetic genes directly and indirectly regulated by FaRIF. **A)** Heatmap displaying the expression of FaRIF-regulated genes related to phenylpropanoid biosynthesis. Significant DEGs are marked with asterisks (***FDR≤0,0005; **FDR≤0,005; *FDR≤0,05). **B)** ChIP-seq peaks showing FaRIF binding sites and input reads over representative homoeologs of direct targets involved in phenylpropanoid biosynthesis. Detailed information is available in Supplementary Data Set S7.

**Supplementary Figure S10.**
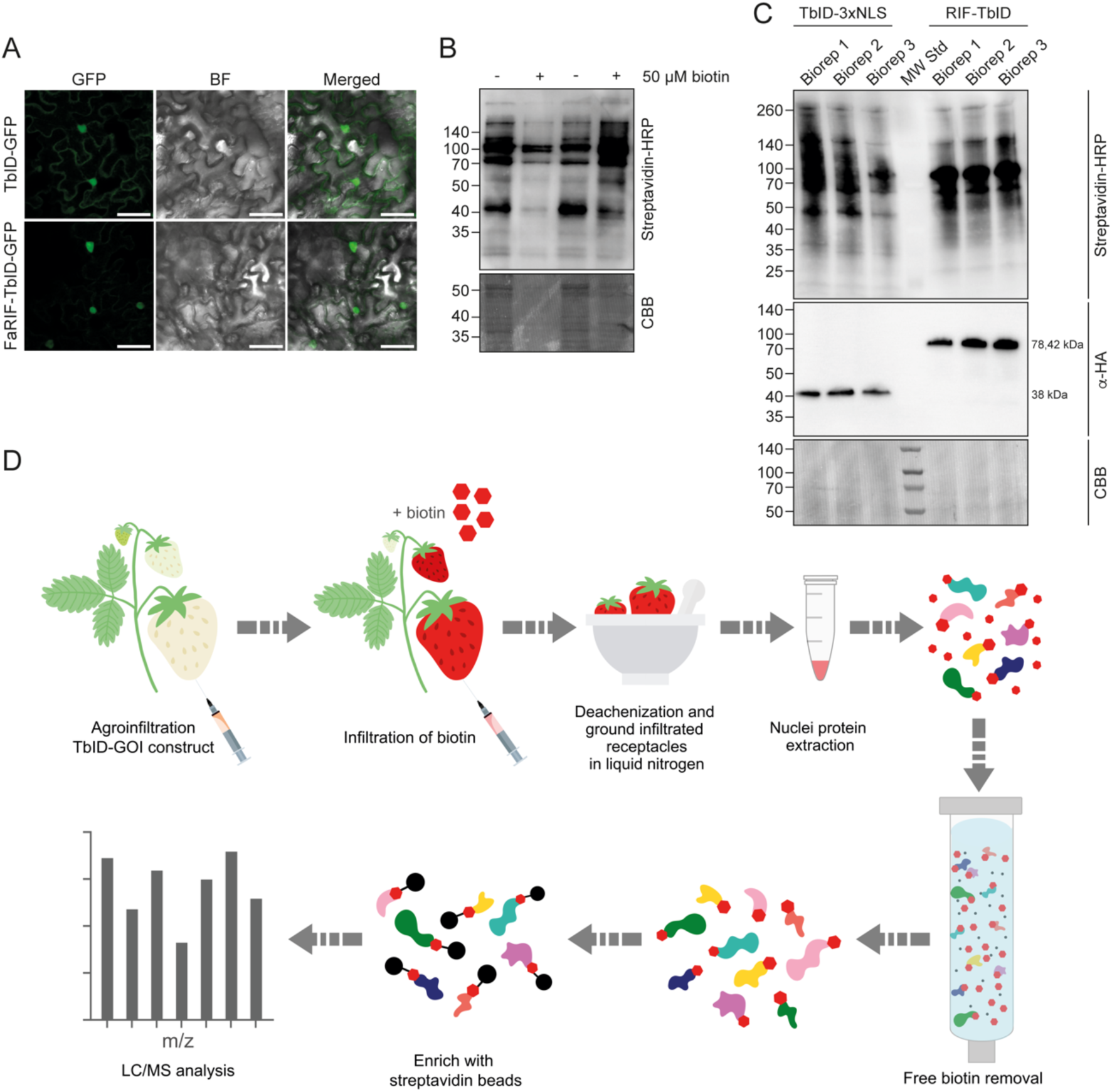
Validation of nuclear localization of FaRIF-TbID construct and biotinylation activity in strawberry fruits. **A)** Subcellular localization of TbID and FaRIF-TbID proteins, both tagged with GFP at the C-terminus in *N. benthamiana* leaves. Images were captured 48h post-infiltration. Scale bars = 50 µm. **B)** Western blot analysis using Streptavidin-HRP of strawberry fruits infiltrated with 50 µM biotin (+) or mock (-) solutions (upper panel). Bottom panel shows Coomassie Brilliant Blue (CBB) staining for loading control. **C)** Western blot analysis of three biological replicates of the FaRIF-TbID fusion protein and the negative control (TbID-3xNLS), used for the TurboID assay and the subsequent GC-MS analysis. The upper panel shows detection with Streptavidin-HRP (upper gel), the middle panel shows detection with αHA antibody, and the lower panel shows CBB staining for loading control. MW Std: Molecular weight standard marker. **D)** Schematic diagram of the TurboID-based proximity labeling pipeline used in this study.

## Supplementary Data Sets

**Supplementary Data Set S1.** Protein sequences of RIF proteins in *F. vesca* (FvRIF) and in *F.* × *ananassa* (FaRIF homoeologs). Gene IDs are included.

**Supplementary Data Set S2.** List of all FaRIF binding sites identified by ChIP-seq in both biological replicates, including those within the region spanning 2 kb upstream to 100 bp downstream of the TSSs.

**Supplementary Data Set S3.** Distribution analysis of FaRIF ChIP-seq peaks (all peaks and those located 2 kbp upstream to 100 bp downstream of TSS) per chromosomes and subgenomes.

**Supplementary Data Set S4.** Transcriptome analysis in control and *35Spro:RIF-* RNAi lines at white and red stages of receptacle ripening. List of DEGs at white and red stages.

**Supplementary Data Set S5.** List of FaRIF direct and indirect targets (DEGs with LogFC≥1 or ≤-1 with and without FaRIF binding sites in their promoter (-2kb to 100 bp TSS) region. Chi-squared analysis in down- and upregulated FaRIF direct target genes.

**Supplementary Data Set S6.** MapMan bins enrichment analysis for DEGs in *FaRIF*-RNAi receptacles at white and red stages, and for FaRIF direct targets.

**Supplementary Data Set S7.** Genes encoding for transcription factors, cell wall and carbohydrate metabolism enzymes, phytohormones biosynthesis and signaling, and phenylpropanoid metabolism enzymes identified among the direct and indirect targets of FaRIF.

**Supplementary Data Set S8**. Proteins, protein groups, and peptides identified in the TurboID proximity labeling assay.

**Supplementary Data Set S9.** List of primers used in this study.

