## Supplementary Data Set S1 for "FaRIF at the Core of Strawberry Fruit Ripening: Deciphering its Targets and Interaction Networks"

```
>FvRIF (FvNAC035)
MESTDSSSGSQPPPQPNLPPGFRFHPTDEELVVHYLKKKASSAPLPVAIIAEVDLYKFD
PWQLPEKATFGEQEWYFFSPRDRKYPNGARPNRAATSGYWKATGTDKPVLSTTDEGGGG
TRKVGVKKALVFYRGKPPKGIKTNWIMHEYRIADNNTSNKPPPGCHDLGNKKNSLRLDD
WVLCRIYKKNNTHRPMDLEDSMDGTMGSSFPLSKLHHLPPKSTTSTYGQFMDNDHNFYD
GMVSSEGINTSASFLPNSALANSSLPLKRELPNLYWNHDPEDEAGPSRRLHMDSSDQST
GNGSIAILLSQLPQTPPPLHQQPMLGTSQLGGDALFRSTQYQLPGNWYS

>FaRIF(3A) (FxaC_9g32650)
MESTDSSSGSQPPPQPNLPPGFRFHPTDEELVVHYLKKKASSAPLPVAIIAEVDLYKFD
PWQLPEKATFGEQEWYFFSPRDRKYPNGARPNRAATSGYWKATGTDKPVLSTTDEGGGG
TRKVGVKKALVFYRGKPPKGIKTNWIMHEYRIADNNTSNKPPPGCHDLGNKKNSLRLDD
WVLCRIYKKNNTHRPMDLEDSMDGTMGSSFPLSKLHHLPPKSTTSTYGQFMDNDHNFYD
GMVSSEGINTSASFLPNSALANSSLPLKRELPNLYWNHDPEDEAGPSRRLHMDSSDQST
GNGSIAILLSQLPQTPPPLHQQPMLGTSQLGGDALFRSTQYQLPGNWYS

>FaRIF(3B) (FxaC_10g22240)
MESTDSSSGSQPPPQPNLPPGFRFHPTDEELVVHYLKKKASSAPLPIAIIAEVDLYKFD
PWQLPEKATFGEQEWYFFSPRDRKYPNGARPNRAATSGYWKATGTDKPVLSSTDEGGGG
TRKVGVKKALVFYRGKPPKGIKTNWIMHEYRLADNNTSNKPPPGCHDLGSKKNSLRLDD
WVLCRIYKKNNTHRPMDQEESMDDMMGSSFPLSKLHRLPPKSTTSTYGQFMDNYHNFYD
GMVSSEGINTSASFLPNSGMANSSLPLKRGLPNLYWNHDQEDEAGLSRRLHLDNSESTG
NGSIATLLSQLPQTPPPLHQQPMLGSQLGGDALFRSTQYQLPGMNWYS

>FaRIF(3C) (FxaC_11g20020)
MESTDSSSGSQQPPPQPNLPPGFRFHPTDEELVVHYLKKKASSAPLPVAIIAEVDLYKF
DPWQLPEKATFGEQEWYFFSPRDRKYPNGARPNRAATSGYWKATGTDKPVLSSTDEGGG
GTRKVGVKKALVFYRGKPPKGIKTNWIMHEYRLADNNTSNKPPPGCHDLGNKKNSLRLD
DWVLCRIYKKNNTHRPMDQEDSMDDMMRSSFPLSNLHRLPPKSTTSTYGQFMDNDHNFY
DGMVSSEGINTSASFLPNSGMANRSLPLKRGLPNLYWNHDQEDEAGLSRRLHLDNSEST
GNGSIATLLSQLPQTPPPLHQQPMLGSQLGGDALFRSTQYQLPGMNWYS

>FaRIF(3D) (FxaC_12g28600)
MHEYRLADNNTSNKPPPGCHDLGNKKNSLRLDDSVLCRIYKKNNTHRLMDQEDSMDDMM
GSSFPLSKLHRLPPKSTTSTYGQFMDNDHNFHDGMEDEAGLSRRLHLDNSESTGNGSIA
TLLSQLPQTPPPLHQQPMLGSQLGGDALFRSTQYQLPGMNWYS
```

**Supplemental Data Set S1.** Protein sequences of RIF proteins in *F. vesca* (FvRIF) and in *F.* × *ananassa* (FaRIF homoeologs). Gene IDs are included.
